# Massively parallel digital transcriptional profiling of single cells

**DOI:** 10.1101/065912

**Authors:** Grace X.Y. Zheng, Jessica M. Terry, Phillip Belgrader, Paul Ryvkin, Zachary W. Bent, Ryan Wilson, Solongo B. Ziraldo, Tobias D. Wheeler, Geoff P. McDermott, Junjie Zhu, Mark T. Gregory, Joe Shuga, Luz Montesclaros, Donald A. Masquelier, Stefanie Y. Nishimura, Michael Schnall-Levin, Paul W Wyatt, Christopher M. Hindson, Rajiv Bharadwaj, Alexander Wong, Kevin D. Ness, Lan W. Beppu, H. Joachim Deeg, Christopher McFarland, Keith R. Loeb, William J. Valente, Nolan G. Ericson, Emily A. Stevens, Jerald P. Radich, Tarjei S. Mikkelsen, Benjamin J. Hindson, Jason H. Bielas

## Abstract

Characterizing the transcriptome of individual cells is fundamental to understanding complex biological systems. We describe a droplet-based system that enables 3′ mRNA counting of up to tens of thousands of single cells per sample. Cell encapsulation in droplets takes place in ∼6 minutes, with ∼50% cell capture efficiency, up to 8 samples at a time. The speed and efficiency allow the processing of precious samples while minimizing stress to cells. To demonstrate the system′s technical performance and its applications, we collected transcriptome data from ∼¼ million single cells across 29 samples. First, we validate the sensitivity of the system and its ability to detect rare populations using cell lines and synthetic RNAs. Then, we profile 68k peripheral blood mononuclear cells (PBMCs) to demonstrate the system′s ability to characterize large immune populations. Finally, we use sequence variation in the transcriptome data to determine host and donor chimerism at single cell resolution in bone marrow mononuclear cells (BMMCs) of transplant patients. This analysis enables characterization of the complex interplay between donor and host cells and monitoring of treatment response. This high-throughput system is robust and enables characterization of diverse biological systems with single cell mRNA analysis.

---

Understanding of biological systems requires the knowledge of their individual components. Single cell RNA-sequencing (scRNA-seq) can be used to dissect transcriptomic heterogeneity that is masked in population-averaged measurements^1,2^. scRNA-seq studies have led to the discovery of novel cell types and provided insights into regulatory networks during development^3^. However, previously described scRNA-seq methods face practical challenges when scaling to tens of thousands or more cells or when it is necessary to capture as many cells as possible from a limited sample^4-9^ (**Supplementary Table 1**). Commercially-available, microfluidic-based approaches have limited throughput^5,6^. Plate-based methods often require time-consuming fluorescence-activated cell sorting (FACS) into many plates that must be processed separately^4,9^. Droplet-based techniques have enabled processing of tens of thousands of cells in a single experiment^7,8^, but current approaches require generation of custom microfluidic devices and reagents.

To overcome these challenges, we developed a droplet-based system that enables 3’ mRNA digital counting of up to tens of thousands of single cells. ∼50% of cells loaded into the system can be captured, and up to 8 samples can be processed in parallel per run. Reverse transcription takes place inside each droplet, and barcoded cDNAs are amplified in bulk. The resulting libraries then undergo standard Illumina short-read sequencing. An analysis pipeline, Cell Ranger, processes the sequencing data and enables automated cell clustering. Here, we first demonstrated comparable sensitivity of the system to existing droplet-based methods by performing scRNA-seq on cell lines and synthetic RNAs. Then, we profiled 68k fresh peripheral blood mononuclear cells (PBMCs) and demonstrated the scRNA-seq platform’s ability to dissect large immune populations. Lastly, we developed a computational method to distinguish donor from host cells in bone marrow transplant samples by genotype. We combined this method with clustering analysis to compare sub-population changes of AML patients. This provided insights into the erythroid lineage in post-transplant samples that have not been possible using morphologic or routine flow cytometry analysis.

## RESULTS

### Droplet-based platform enables barcoding of tens of thousands of cells

The scRNA-seq microfluidics platform builds on the GemCode® technology, which has been used for genome haplotyping, structural variant analysis and *de novo* assembly of a human genome^10-12^. The core of the technology is a Gel bead in Emulsion (GEM). GEM generation takes place in an 8-channel microfluidic chip that encapsulates single gel beads at ∼80% fill rate (**Fig. 1a-c**). Each gel bead is functionalized with barcoded oligonucleotides that consist of: i) sequencing adapters and primers, ii) a 14bp barcode drawn from ∼750,000 designed sequences to index GEMs, iii) a 10bp randomer to index molecules (unique molecular identifier, UMI), and iv) an anchored 30bp oligo-dT to prime poly-adenylated RNA transcripts (**Fig. 1d**). Within each microfluidic channel, ∼100,000 GEMs are formed per ∼6-min run, encapsulating thousands of cells in GEMs. Cells are loaded at a limiting dilution to minimize co-occurrence of multiple cells in the same GEM.

**Figure 1.**
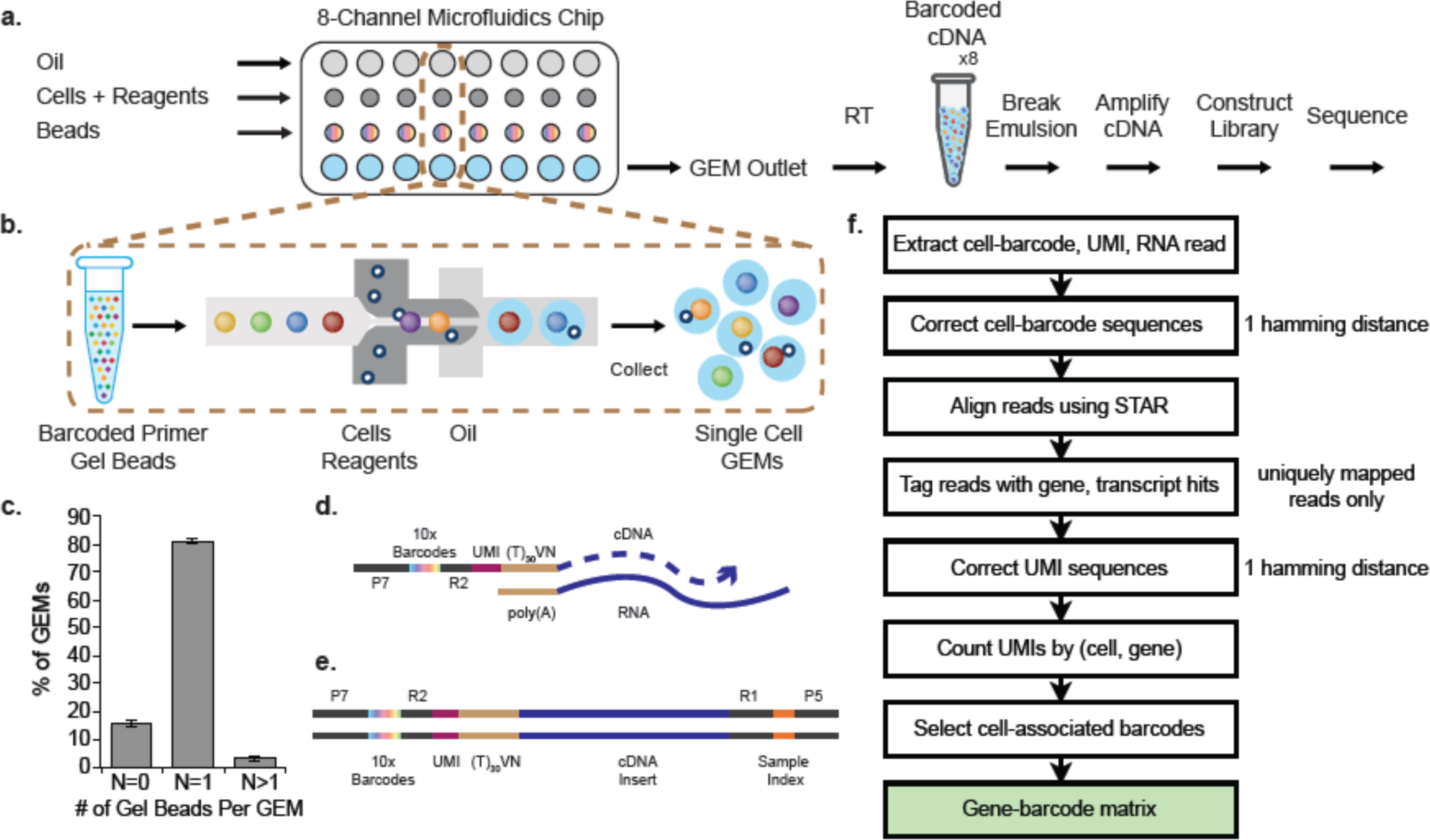
GemCode single cell technology enables 3’ profiling of RNAs from thousands of single cells simultaneously. **(a)** scRNA-seq workflow on GemCode technology platform. Cells were combined with reagents in one channel of a microfluidic chip, and gel beads from another channel to form GEMs. RT takes place inside each GEM, after which cDNAs are pooled for amplification and library construction in bulk. **(b)** Gel beads loaded with primers and barcoded oligonucleotides are first mixed with cells and reagents, and subsequently mixed with oil-surfactant solution at a microfluidic junction. Single cell GEMs are collected in the GEM Outlet. **(c)** % GEMs containing 0 gel bead (N = 0), 1 gel bead (N = 1) and >1 gel bead (N = 2). Data include 5 independent runs from multiple chip and gel bead lots over >70k GEMs for each run, n = 5, mean ± s.e.m. **(d)** Gel beads contain barcoded oligonucleotides consisting of Illumina adapters, 10x barcodes, UMIs and oligo dTs, which prime reverse transcription of poly-adenylated RNAs. **(e)** Finished library molecules consist of Illumina adapters and sample indices, allowing pooling and sequencing of multiple libraries on a next generation short read sequencer. **(f)** Cell Ranger pipeline workflow. Gene-barcode matrix (highlighted in green) is an output of the pipeline.

Cell lysis begins immediately after encapsulation. Gel beads automatically dissolve to release their oligonucleotides for reverse transcription of poly-adenylated RNAs. Each cDNA molecule contains a UMI and shared barcode per GEM, and ends with a template switching oligo at the 3’ end (**Fig. 1e**). Next, the emulsion is broken and barcoded cDNA is pooled for PCR amplification, using primers complementary to the switch oligos and sequencing adapters. Finally, amplified cDNAs are sheared, and adapter and sample indices are incorporated into finished libraries which are compatible with next-generation short-read sequencing. Read1 contains the cDNA insert while Read2 captures the UMI. Index reads, I5 and I7, contain the sample indices and cell barcodes respectively. This streamlined approach enables parallel capture of thousands of cells in each of the 8 channels for scRNA-seq analysis.

### Technical demonstration with cell lines and synthetic RNAs

To assess the technical performance of our system, we loaded a mixture of ∼1,200 human (293T) and ∼1,200 mouse (3T3) cells and sequenced the library on the Illumina NextSeq 500 to yield ∼100k reads/cell. Sequencing data were processed by Cell Ranger (Online Methods, **Fig. 1f**). Briefly, 98-nt of Read1s were aligned against the union of human (hg19) and mouse (mm10) genomes with STAR. Barcodes and UMIs were filtered and corrected. PCR duplicates were marked using the barcode, UMI and gene ID. Only confidently mapped, non-PCR duplicates with valid barcodes and UMIs were used to generate the gene-barcode matrix for further analysis. 38% and 33% of reads mapped to human and mouse exonic regions, respectively and <6% of reads mapped to intronic regions (**Supplementary Table 2**). The high mapping rate is comparable to previously reported scRNA-seq systems^4-9^.

Based on the distribution of total UMI counts for each barcode (Online Methods), we estimated that 1,012 GEMs contained cells, of which 482 and 538 contained reads that mapped primarily to the human and mouse transcriptome, respectively (and will be referred to as human and mouse GEMs) (**Fig. 2a**). >83% of UMI counts were associated with cell barcodes, indicating low background of cell-free RNA. Eight cell-containing GEMs had a substantial fraction of human and mouse UMI counts (the UMI count is >1% of each species’ UMI count distribution), yielding an inferred multiplet rate (rate of GEMs containing >1 cell) of 1.6% (Online Methods, **Fig. 2a**). A cell titration experiment across six different cell loads showed a linear relationship between the multiplet rate and the number of recovered cells ranging from 1,200 to 9,500 (**Supplementary Fig. 1a**). The multiplet rate and trend are consistent with Poisson loading of cells, and have been validated by independent imaging experiments (Online Methods, **Supplementary Fig. 1b**). In addition, we observed ∼50% cell capture rate, which is the ratio of the number of cells detected by sequencing and the number of cells loaded. The capture rate is consistent across four types of cells with cell loads ranging from ∼1,000 to ∼23,000 (**Supplementary Table 3**), a key improvement over some existing scRNA-seq methods (**Supplementary Table 1**). Lastly, the mean fraction of UMI counts from the other species was 0.9% in both human and mouse GEMs, indicating a low level of cross-talk between cell barcodes (Online Methods). This, coupled with the low multiplet rate and high cell capture rate, is particularly important for samples with limited cell input and for the detection of rare cells.

**Figure 2.**
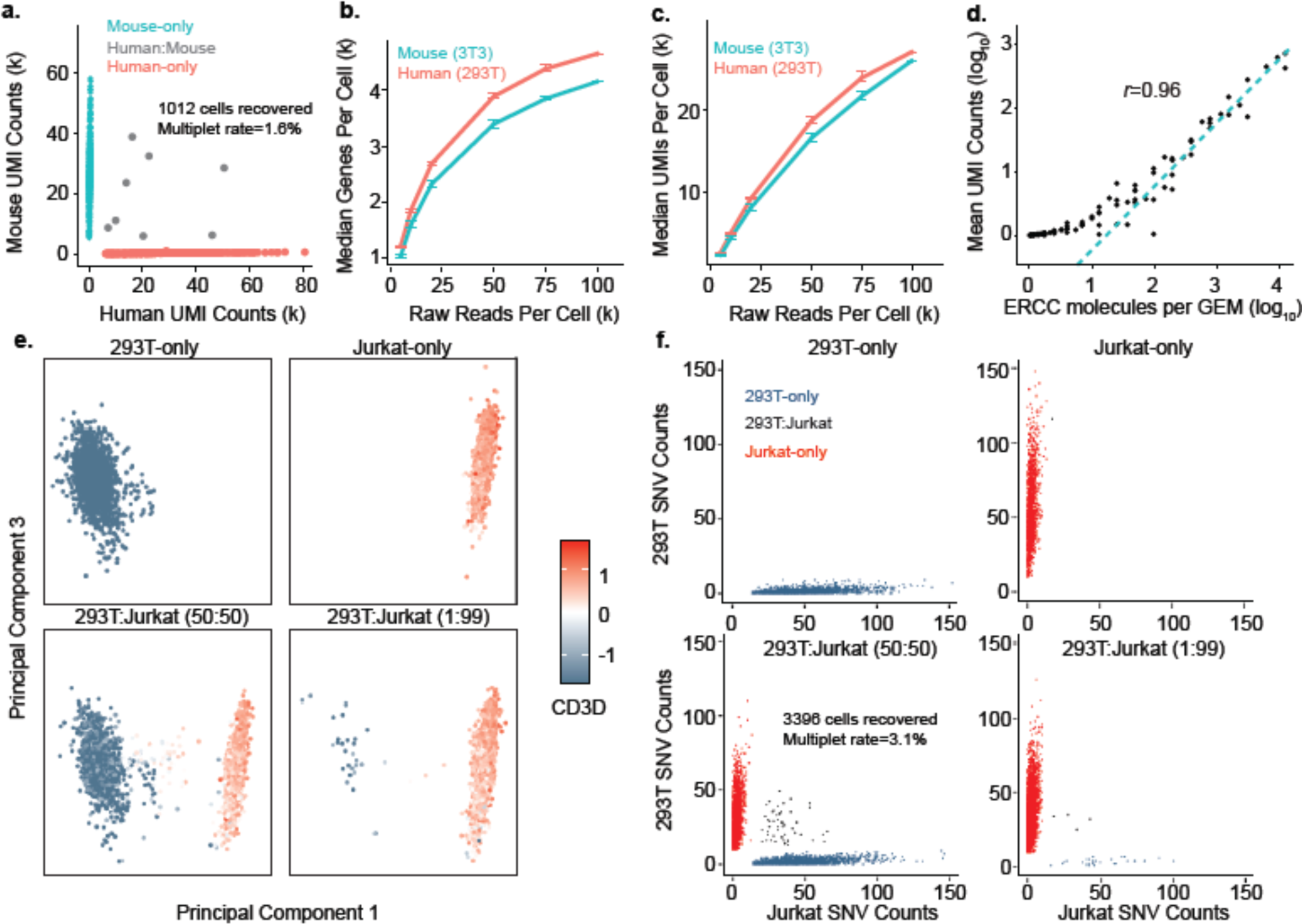
Demonstration of technical performance of GemCode single cell technology with cell lines and ERCC. **(a)** Scatter plot of human and mouse UMI counts detected in a mixture of 293T and 3T3 cells. Cell barcodes containing primarily mouse reads are colored in cyan and termed “Mouse-only”; cell barcodes with primarily human reads are colored in red and termed “Human-only”; and cell barcodes with significant mouse and human reads are colored in grey and termed “Human:Mouse”. A multiplet rate of 1.5% was inferred. Median number of genes **(b)** and UMI counts **(c)** detected per cell in a mixture of 293T (red) and 3T3 (cyan) cells at different raw reads per cell. Data from 3 independent experiments were included, mean ± s.e.m. **(d)** Mean observed UMI counts for each ERCC molecule is compared to expected number of ERCC molecules per GEM. A straight line was fitted to summarize the relationship. **(e)** Principal component (PC) analysis was performed on normalized scRNA-seq data of Jurkat and 293T cells mixed at 4 different ratios (100% 293T, 100% Jurkat, 50:50 293T:Jurkat and 1:99 293T and Jurkat). PC1 and PC3 are plotted, and each cell is colored by the normalized expression of CD3D. **(e)** SNVs analysis was performed, and 293T-enriched SNVs and Jurkat-enriched SNVs were plotted for each sample. A 3.1% multiplet rate was inferred from the 50:50 293T:Jurkat sample.

The sensitivity of scRNA-seq methods is critical to many applications. At 100k reads/cell, we detected a median of ∼4,500 genes and 27,000 transcripts (UMI counts) in each human and mouse cell, indicating comparable sensitivity to other droplet-based platforms^7,8^ (**Fig. 2b, c**). UMI counts showed a standard deviation of ∼43% of the mean (CV) in human cells, and ∼33% of the mean in mouse cells, where the trend is consistent in four independent human and mouse mixture experiments (**Supplementary Fig. 1c,d**). Genes of different GC composition and length show similar UMI count distributions, suggesting low transcript bias (**Supplementary Fig. 1e-h**).

We also directly measured cDNA conversion rate by loading External RNA Controls Consortium (ERCC) synthetic RNAs into GEMs in place of cells. We found that mean UMI counts from sequencing was highly correlated (r=0.96) with molecule counts calculated from the loading concentration of ERCC (**Fig. 2d**, **Supplementary Fig. 2a**). Furthermore, we inferred 6.7-8.1% efficiency from both ERCC RNA Spike-in Mix1 and Mix2 in a 1:50 dilution (**Supplementary Fig. 2b**), with minimal evidence of GC bias, and limited bias for transcripts longer than 500-nt (**Supplementary Fig. 2c, d**). Additionally, we estimated the conversion rate of cell transcripts in Jurkat cells by ddPCR^13^. The amount of cDNA of 8 genes obtained from single cells after reverse transcription in GEMs was compared to the expected RNA inferred from bulk profiling. The conversion rates among 8 genes were between 2.5 and 25.5%, which is consistent with ERCC data (Online Methods, **Supplementary Fig. 2e**).

The ERCC experiments also allowed us to estimate the relative proportion of biological and technical variation. Since ERCCs are in solution, they do not introduce biological variation related to differences in cell size, RNA content or transcriptional activity. Thus, technical variation is the only source of variation. When the ERCCs are dilute (UMI counts are small), sampling noise dominates; when the UMI counts increase, technical variations become dominant^14^ (**Supplementary Fig. 2f**). These variations include variation in droplet size, variation in concentration of RT reagents in the droplets, variation in the concentration of sample in the droplets, and variation in RT and/or PCR efficiency of the distinct gel bead barcode sequences. The squared coefficient of variation (CV^2^) is ∼7% among all the ERCC experiments. In comparison, CV^2^ in samples of mouse and human cells is ∼11-19% (**Supplementary Fig. 1d**), suggesting that technical variance accounts for ∼50% of total variance, consistent with the observations from Klein *et al*^8^.

### Detection of individual populations in *in-vitro* mixed samples

We tested the ability of the system to accurately detect heterogeneous populations by mixing two cell lines, 293T and Jurkat cells at different ratios (**Supplementary Table 2**). We performed principal component analysis (PCA) on UMI counts from all detected genes after pooling all the samples (**Supplementary Fig. 3a**). In the sample where an equal number of 293T and Jurkat cells was mixed, principal component (PC) 1 separated cells into two clusters of equal size (**Fig. 2e**, **Supplementary Fig. 4a**, **Supplementary Table 4**). Based on expression of cell type specific markers, we infer that one cluster corresponds to Jurkat cells (preferentially expressing CD3D), and the other corresponds to 293T cells (preferentially expressing XIST, as 293T is a female cell line, and Jurkat is a male cell line) (**Fig. 2e**, **Supplementary Fig. 4b**). Points located between the two clusters are likely multiplets, as they expressed both CD3D and XIST (**Fig. 2e**, **Supplementary Fig. 4b)**. In contrast, PC1 did not separate cells into two clusters in the 293T-only and the Jurkat-only samples (**Fig. 2e**). Furthermore, in the sample with 1% 293T and 99% Jurkat cells, the number of cells in each of the two clusters were observed at the expected ratio (**Fig. 2e**, **Supplementary Fig. 4a**, **Supplementary Fig. 4b**). A similar trend was observed for 12 independent samples where 293T and Jurkat cells were mixed at 5 different proportions, demonstrating the system’s ability to perform unbiased detection of rare single cells (**Supplementary Fig. 4a**).

Our scRNA-seq data not only provides a digital transcript count, it also provides ∼250nt sequence for each cDNA that could be used for Single Nucleotide Variant (SNV) detection. On average, there are ∼350 SNVs detected in each 293T or Jurkat cell (**Supplementary Fig. 4c**, **Supplementary Table 5**), and we investigated whether the SNVs could be used independently to distinguish cells in the mixture. We selected a set of high quality SNVs that were only observed in 293T or Jurkat cells, but not both (Online Methods). We then scored cells in the mixed samples based on the number of 293T or Jurkat-enriched SNVs (Online Methods). In the 1:1 mixed sample, 45% 293T cells primarily (96%) harbored 293T-enriched SNVs, whereas 50% Jurkat cells primarily (94%) harbored Jurkat-enriched SNVs (**Fig. 2f**, **Supplementary Table 6**). Jurkat and 293T cells inferred from marker-based analysis is 99% consistent with SNV-based assignment. We observed a multiplet rate of ∼3%, accounting for multiplets from Jurkats:293Ts as well as Jurkats:Jurkats and 293Ts:293Ts. The multiplet rate is consistent with that predicted from the human and mouse mixing experiment, when ∼3000 cells were recovered (**Supplementary Fig. 1a**). Our result demonstrates that SNVs detected from scRNA-seq data can be used to classify individual cells.

### Subpopulation discovery from a large immune population

The GemCode single cell technology can also be used for scRNA-seq of primary cells. To study immune populations within PBMCs, we obtained fresh PBMCs from a healthy donor (Donor A). ∼8-9k cells were captured from each of 8 channels and pooled to obtain ∼68k cells. Data from multiple sequencing runs were merged using the Cell Ranger pipeline. At ∼20k reads/cell, the median number of genes and UMI counts detected per cell was approximately 525 and 1,300, respectively (**Fig. 3a**, **Supplementary Fig. 5a**). The UMI count is roughly 10% of that from 293T and 3T3 samples at ∼20k reads/cell, likely reflecting the differences in cells’ RNA content (∼1pg RNA/cell in PBMCs vs. ∼15pg RNA/cell in 293T and 3T3 cells) (**Supplementary Fig. 5a, b**).

**Figure 3.**
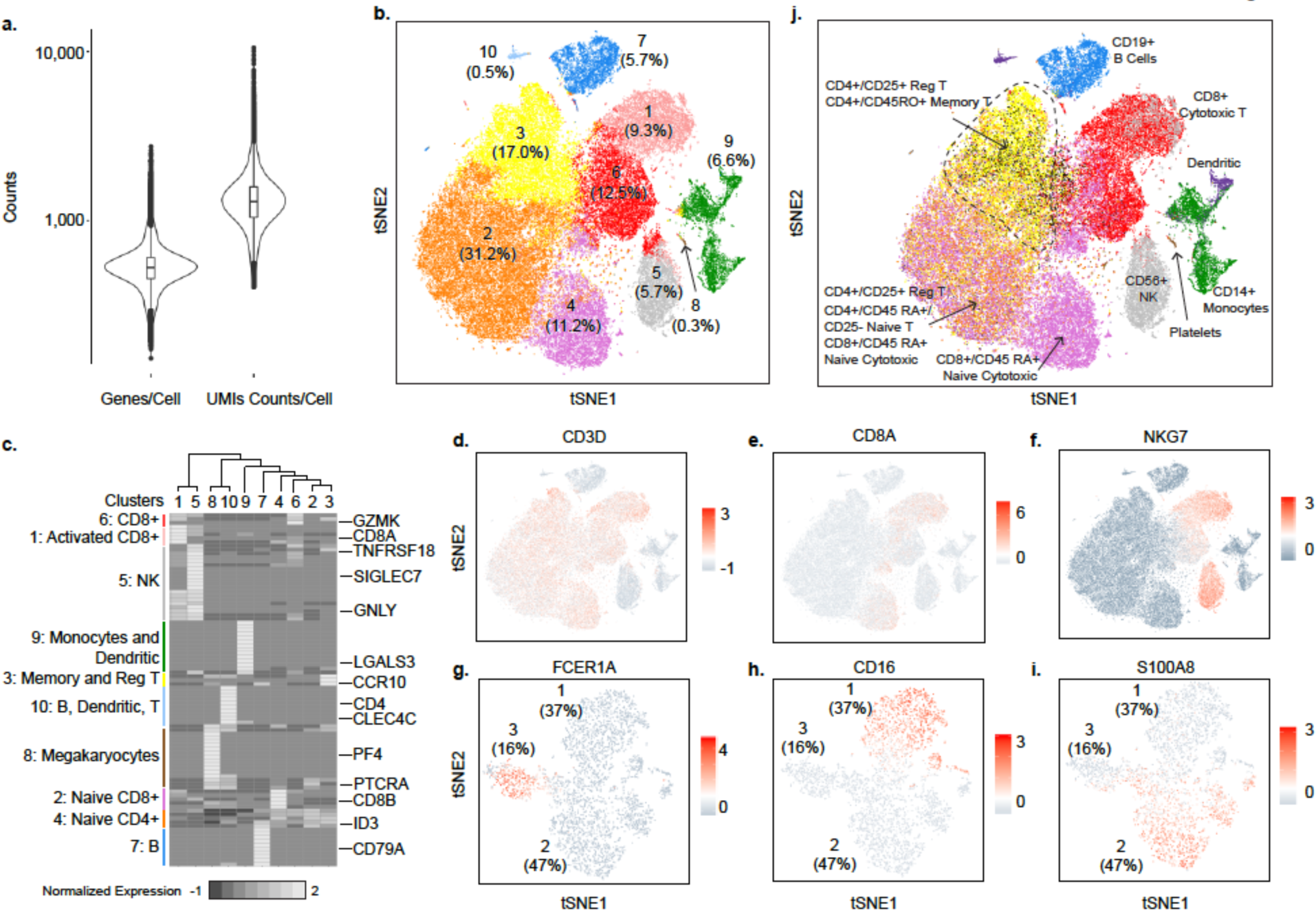
Distinct populations can be detected in fresh 68k PBMCs. **(a)** Distribution of number of genes (left) and UMI counts (right) detected per 68k PBMCs. **(b)** tSNE projection of 68k PBMCs, where each cell is grouped into one of the 10 clusters (distinguished by their colors) Cluster number is indicated, with the percentage of cells in each cluster noted in parentheses. **(c)** Normalized expression (centered) of the top variable genes (rows) from each of 10 clusters (columns) is shown in a heatmap. Numbers at the top indicate cluster number in **b**, with connecting lines indicating the hierarchical relationship between clusters. Representative markers from each cluster are shown on the right, and an inferred cluster assignment is shown on the left. **(d-i)** tSNE projection of 68k PBMCs, with each cell colored based on their normalized expression of CD3D, CD8A, NKG7, FCER1A, CD16 and S100A8. UMI normalization was performed by first dividing UMI counts by the total UMI counts in each cell, followed by multiplication with the median of the total UMI counts across cells. Then we took the natural log of the UMI counts. Finally, each gene was normalized such that the mean signal for each gene is 0, and standard deviation is 1. **(j)** tSNE projection of 68k PBMCs, with each cell colored based on their correlation-based assignment to a purified sub-population of PBMCs. Sub-clusters within T cells are marked by dashed polygons.

We performed clustering analysis to examine cellular heterogeneity among PBMCs. We applied PCA on the top 1000 variable genes ranked by their normalized dispersion, following a similar approach to Macosko *et al.*^7^ (**Supplementary Fig. 3b**, **5c**, Online Methods). K-means^15^ clustering on the first 50 PCs identified 10 distinct cell clusters, which were visualized in two dimensional projection of t-Distributed Stochastic Neighbor Embedding (tSNE)^16^ (Online Methods, **Fig. 3b**, **Supplementary Fig. 5d**). To identify cluster-specific genes, we calculated the expression difference of each gene between that cluster and average of the rest of clusters. Examination of the top cluster-specific genes revealed major subtypes of PBMCs at expected ratios^17^: >80% T cells (enrichment of CD3D, part of the T cell receptor complex, in clusters 1-3, and 6), ∼6% NK cells (enrichment of NKG7^18^ in cluster 5), ∼6% B cells (enrichment of CD79A^19^ in cluster 7) and ∼7% myeloid cells (enrichment of S100A8 and S100A9^20^ in cluster 9 (Online Methods, **Fig. 3b-f**, **Supplementary Fig. 5e**, **Supplementary Table 7**). Finer substructures were detected within the T cell cluster; clusters 1, 4 and 6 are CD8+ cytotoxic T cells, whereas clusters 2 and 3 are CD4+ T cells (**Fig. 3e**, **Supplementary Fig. 5f**). The enrichment of NKG7 on cluster 1 cells implies a cluster of activated cytotoxic T cells^21^ (**Fig. 3f**). Cells in Cluster 3 showed high expression of CCR10 and TNFRSF18, markers for memory T cells^22^ and regulatory T cells^23^ respectively, and likely consisted of a mixture of memory and regulatory T cells (**Fig. 3c**, **Supplementary Fig. 5g**). The presence of ID3, which is important in maintaining a naïve T cell state^24^, suggests that cluster 2 represents naïve CD8 T cells whereas cluster 4 represents naïve CD4 T cells (**Fig. 3c**). To identify sub-populations within the myeloid population, we further applied k-means clustering on the first 50 PCs of cluster 9 cells. At least 3 populations were evident: dendritic cells (characterized by presence of FCER1A^25^), CD16+ monocytes, and CD16-/low monocytes^26^ (**Fig. 3g-i**, **Supplementary Table 7**). Overall, these results demonstrate that our scRNA-seq method can detect all major subpopulations expected to be present a PBMC sample.

Our analysis also revealed some minor cell clusters, such as cluster 8 (0.3%) and cluster 10 (0.5%) (**Fig. 3b**). Cluster 8 showed preferential expression of megakaryocyte markers, such as PF4, suggesting that it represents a cluster of megakaryocytes (**Fig. 3b-c**, **Supplementary Fig. 5h**). Cells in cluster 10 express markers of B, T and dendritic cells, suggesting a likely cluster of multiplets (**Fig. 3b, c**). The size of the cluster suggests the multiplets comprised mostly of B:dendritic and B:T:dendritic cells (Online Methods). With ∼9k cells recovered per channel, we expect a ∼9% multiplet rate and that the majority of multiplets would only contain T cells. More sophisticated methods will be required to detect multiplets from identical or highly similar cell types.

To further characterize the heterogeneity among 68k PBMCs, we generated reference transcriptome profiles through scRNA-seq of 10 bead-enriched subpopulations of PBMCs from Donor A (**Supplementary Fig. 6-7, Supplementary Table 8**). Clustering analysis revealed a lack of sub-structure in most samples, consistent with the samples being homogenous populations, and in agreement with FACS analysis (Online Methods, **Supplementary Fig. 6-7**). However, substructures were observed in CD34+ and CD14+ monocyte samples (Online Methods, **Supplementary Fig. 7b, j**). In the CD34+ sample, ∼70% cell clusters show expression of CD34 (**Supplementary Fig. 7j**). In the CD14+ sample, the minor population showed marker expression for dendritic cells (e.g. CLEC9A), providing another reference transcriptome to classify the 68k PBMCs (**Supplementary Fig. 7b**). This result also demonstrates the power of scRNA-seq in selecting appropriate cells for further analysis.

We classified 68k PBMCs based on their best match to the average expression profile of 11 reference transcriptomes (Online Methods, **Fig. 3j**). Cell classification was largely consistent with previously described marker-based classification, although the boundaries among some of the T cell sub-populations were blurred. Namely, part of the inferred CD4+ naïve T population was classified as CD8+ T cells. We have also tried to cluster the 68k PBMC data with Seurat^27^ (Online Methods). While it was able to distinguish inferred CD4+ naïve from inferred CD8+ naïve T cells, it was not able to cleanly separate out inferred activated cytotoxic T cells from inferred NK cells (**Supplementary Fig. 5i**). Such populations have overlapping functions, making separation at the transcriptome level particularly difficult and even unexpected. However, the complementary results of Seurat’s and our analysis suggest that more sophisticated clustering and classification methods can help address these problems.

### Single cell RNA profiling of cryopreserved PBMCs

In order to analyze repository specimens for clinical research, we tested GemCode technology on cryopreserved cells. We froze the remaining fresh PBMCs from Donor A, and made a scRNA-seq library from gently thawed cells a week later where ∼3k cells were recovered (Online Methods). The two datasets (fresh and frozen) showed a high similarity between their average gene expression (r=0.97, Online Methods, **Supplementary Fig. 8a**). 57 genes showed 2-fold upregulation in the frozen sample, with ∼50% being ribosomal protein genes, and the rest not enriched in any pathways (**Supplementary Table 9**). In addition, the number of genes and UMI counts detected from fresh and frozen PBMCs was very similar (*p* = 0.8 and 0.1, respectively), suggesting that the conversion efficiency of the system is not compromised when profiling frozen cells (**Supplementary Fig. 8b**). Furthermore, subpopulations were detected from frozen PBMCs at a similar proportion to that of fresh PBMCs, demonstrating the applicability of our method on frozen samples (Online Methods, **Supplementary Fig. 8c**).

### A genotype-based method to delineate individual populations from a mixed sample

Next, we applied the GemCode technology to study host and donor cell chimerism in an allogeneic hematopoietic stem cell transplant (HSCT) setting. Following a stem cell transplant it is important to monitor the proportion of donor and host cells in major cell lineages to ensure complete engraftment and as a sensitive means of detecting impending relapse. Traditionally, the amount of host and donor chimerism is measured from flow sorted cell populations by PCR assays with a panel of SNV-specific primers. This procedure is labor intensive and is limited to a few major lineage populations. Here, we present a simple method to determine host and donor chimerism at single cell resolution that enables extensive characterization of cell subtypes and donor/host genotypes by integrating scRNA-seq with de novo SNV calling.

While previous studies have used existing SNVs from DNA sequencing or large scale copy number changes (CNV) in the transcriptome data to distinguish cells by genotype^28-31^, these methods cannot be applied to transplant samples where donor and host genotype is not known *a priori*, and when donor and host are closely matched in genotype. To this end, we first developed a method to infer the relative presence of host and donor genotypes in a mixed population based on SNVs directly predicted from the transcriptome data. The method identifies SNVs and infers a genotype at each SNV. It then classifies cells based on their genotypes across all SNVs (Online Methods).

To evaluate the technical performance of this method, we generated scRNA-seq libraries from PBMCs of two healthy donors B and C, with ∼8k cells captured for each sample (**Supplementary Table 2**). We first performed *in silico* mixing of PBMCs B and C at 12 mixing ratios ranging from 0 to 50%. Only confidently mapped reads from samples B and C were used, and a total of 6000 cells were selected (Online Methods). There were ∼15k reads/cell, with ∼50 filtered SNVs per cell (Online Methods, **Supplementary Fig. 9a, b**, **Supplementary Tables 2, 5**). We then classified the cells based on variants detected from the mixed transcriptome. Sensitivity and positive predictive value (PPV) were calculated by comparing the predicted call of each cell against its true labeling. Our method was able to identify minor genotypes as low as 3% at >95% sensitivity and PPV (**Fig. 4a, b**). A minor population could not be detected when the mixed ratio was below 3% (**Fig. 4c**). The accuracy of this method is affected by the number of observed SNVs per cell, which is dependent on cell types, diversity between subjects, and variant calling sensitivity. Nevertheless, the base error rate and variant calling errors have a limited effect on the accuracy of the method, as the method uses all instead of a small subset of SNVs (**Supplementary Fig. 9c**).

**Figure 4.**
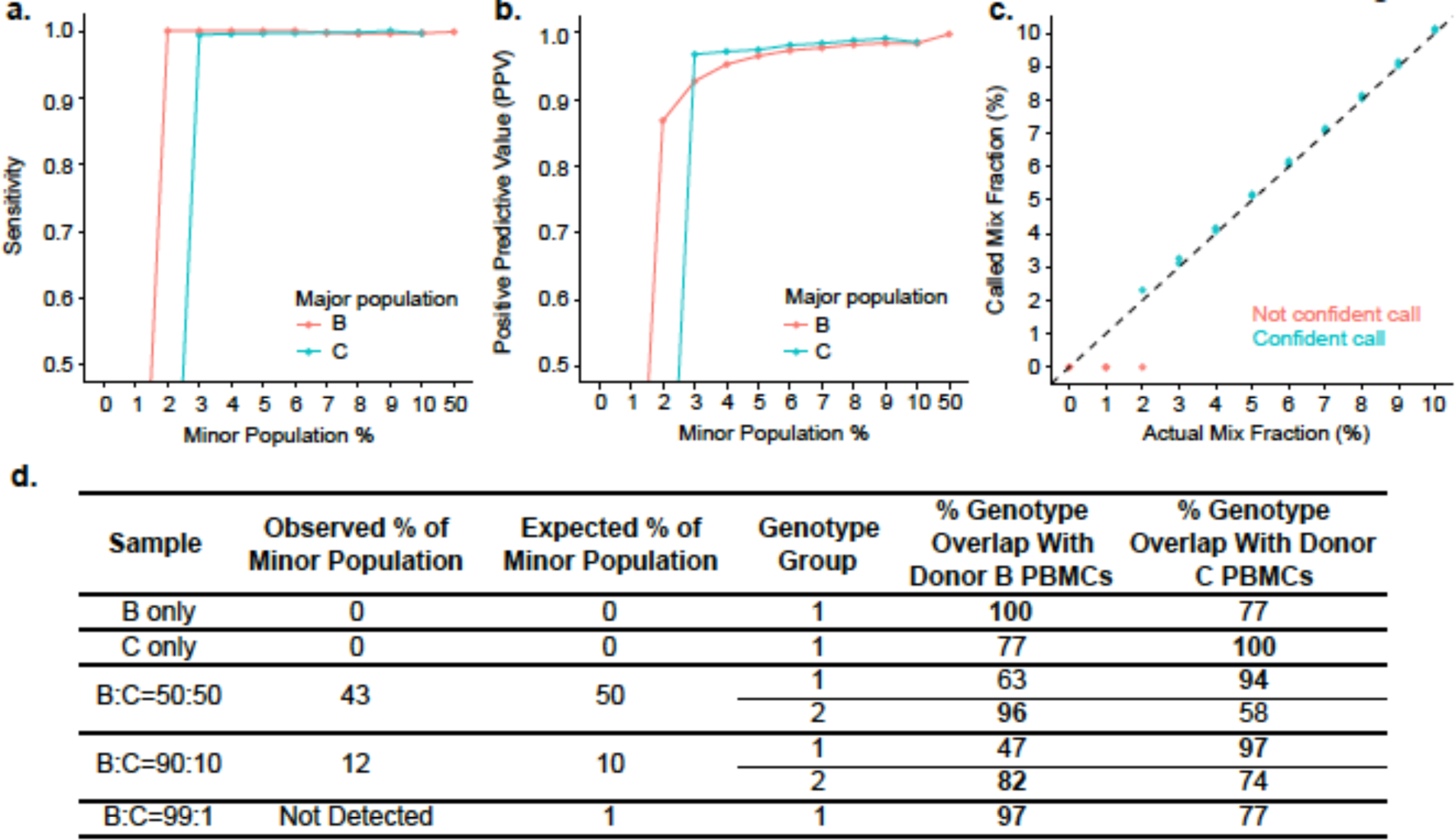
Genotype analysis of *in silico* and *in vitro* mixing of PBMCs. **(a)** Sensitivity vs. % minor population, where sensitivity is evaluated against the true labeling of *in silico* mixed PBMCs from Donors B and C. Red line indicates that the major population comes from Donor B PBMCs. Blue line indicates that the major population comes from Donor C PBMCs. **(b)** PPV vs. % minor population, where PPV is evaluated against the true labeling of in silico mixed PBMCs from Donors B and C. Red line indicates that the major population comes from Donor B cells. Blue line indicates that the major population comes from Donor C cells. **(c)** Called mix fraction vs. actual mix fraction in *in silico* mixing of PBMCs from Donors B and C. 50% actual mix fraction is correctly called, but omitted from the plot so that the rest of the ratios can be clearly displayed. **(d)** Genotype comparison of predicted genotype groups to purified populations.

We further validated the performance of the method in experiments where PBMCs from donors B and C were mixed at three ratios, 50:50, 90:10 and 99:1, prior to scRNA-seq. In the 50:50 mixture sample, cells from donors B and C were almost indistinguishable by RNA expression (**Supplementary Fig. 9d, e**). However, they can be separated by their genotype at the correct proportion (**Fig. 4d**). In addition, the genotype overlap between genotype group 1 and Donor C is 94%, whereas the overlap between genotype group 1 and Donor B is only 63%, both within the range of positive and negative controls, suggesting that group 1 comes from Donor C (Online Methods, **Fig. 4d**). Similarly, genotype group 2 was inferred to be from Donor B (Online Methods, **Fig. 4d**). The proportions of the minor genotype were accurately predicted at the 90:10 mixing ratio. Consistent with the *in silico* mixing results, the minor population could not be detected when B and C were mixed at 99:1 ratio (**Fig. 4d**).

### Single cell analysis of transplant bone marrow samples

Single cell RNA-seq libraries were generated from cryopreserved BMMC samples obtained from two patients before and after undergoing HSCT for acute myeloid leukemia (AML) (AML027 and AML035) (**Supplemental Table 2**). Since HSCT samples are fragile, cells were carefully washed in PBS with 20% FBS before loading them into chips. Relative to BMMCs from 2 healthy controls, we found 3-5 times as many median number of UMI counts per cell in AML samples at ∼15k reads/cell, suggesting their vastly abnormal transcriptional programs (**Supplementary Fig. 10a**). ∼35 and 60 SNVs/cell were detected from AML027 and AML035 pre-transplant samples respectively (**Supplementary Table 5**, **Supplementary Fig. 10b, c**). Our SNV analysis detected the presence of two genotypes in the post-transplant sample of AML027, one at 13.8%, and one at 86.2% (**Fig. 5a**). As expected, there was no evidence of multiple genotype groups in the pre-transplant host sample. We compared the major and minor inferred genotypes present in the post-transplant sample to the genotype found in the host cells. The major inferred genotype in the post-transplant sample was 97% similar to that inferred from the host sample, while the minor inferred genotype was only 52% similar to that of the host sample (**Fig. 5a**). The observed range of genotype overlap between the same individuals is ∼98% (Online Methods), indicating errors in the genotypes inferred from individual SNVs. 97% is within the observed range, and this results suggests that the post-transplant sample consists mainly (86.2%) of host cells. This observation is consistent with the clinical chimerism assay, which demonstrated only 12% donor in the post-transplant sample. In contrast, SNV analysis on the post-HSCT sample from AML035 did not detect the presence of 2 genotype groups. The sample only shared 78% similarity with AML035 host cells, suggesting that the post-HSCT sample was all donor-derived (**Fig. 5a**). This finding was validated by the independent clinical chimerism assay.

**Figure 5.**
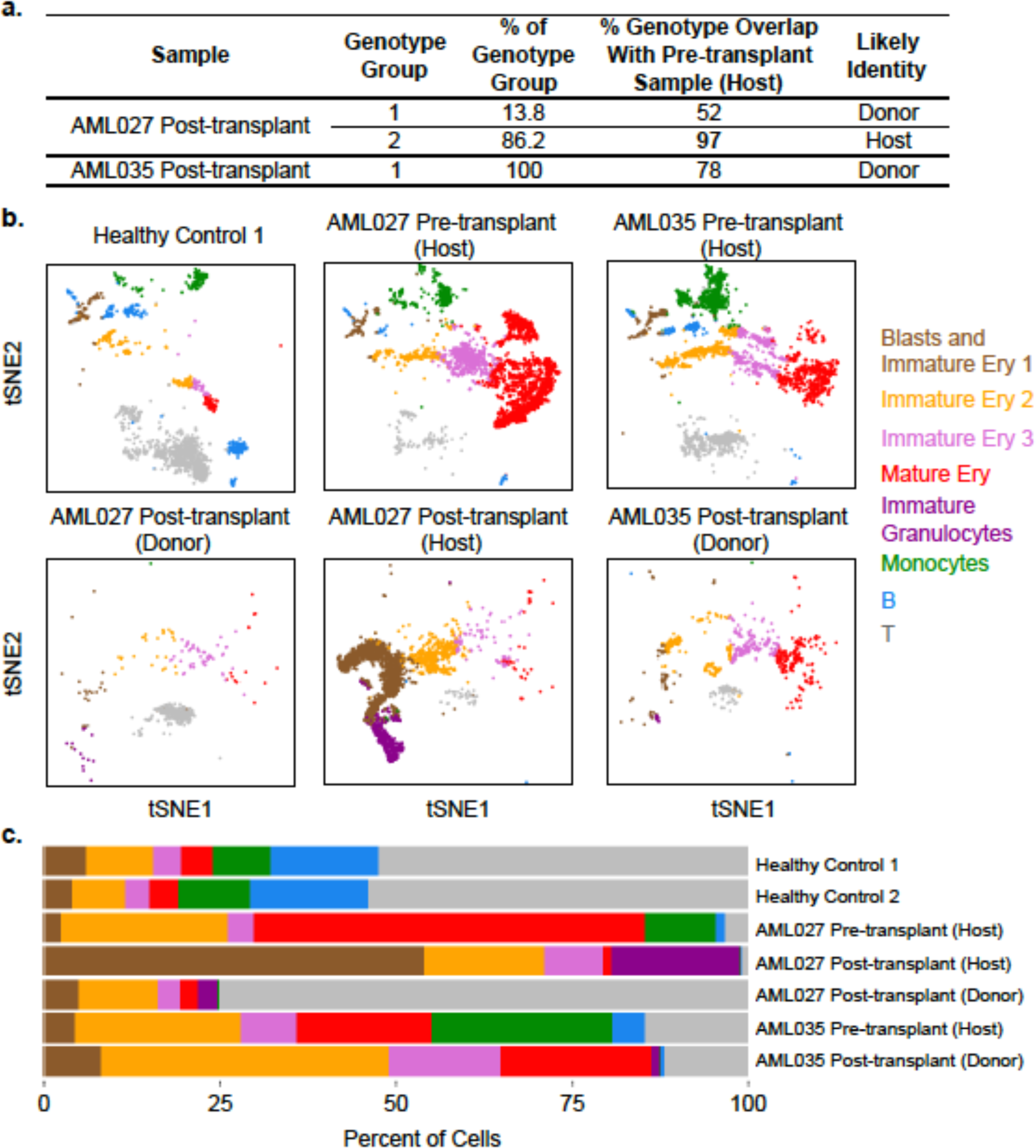
Genotype and single cell expression analysis of transplant BMMCs. **(a)** Predicted genotype groups and their genotype overlap with pre-transplant samples. **(b)** tSNE projection of scRNA-seq data from a healthy control, AML027 pre- and post-transplant samples (post-transplant sample is separated into host and donor), and AML035 pre- and post-transplant samples. tSNE projection was also performed on a 2^nd^ healthy control, but the plot is not included here as it is very similar to that of the first healthy control. Each cell is colored by their classification, which is labeled next to the cell clusters. **(c)** Proportion of sub-populations in each sample.

SNV and scRNA-seq analyses enable subpopulation comparison between individuals within and across multiple samples. We applied these analyses on BMMC scRNA-seq data from healthy controls and AML patients (Online Methods), and observed subpopulation differences in AML patients after HSCT. First, while T cells dominate the healthy BMMCs and donor cells of the AML027 post-transplant sample as expected, erythroids constitute the largest population among AML samples (**Fig. 5b**). Different sets of progenitor and differentiation markers (e.g. CD34, GATA1, CD71 and HBA1) were detected among the erythroids, indicating populations at various stages of erythroid development (Online Methods, **Supplementary Fig. 10d-f**). AML027 showed the highest level of erythroid cells (>80%, consist of mostly mature erythroids) before transplant, consistent with the erythroleukemia diagnosis of AML027 (**Fig. 5c**). In contrast, after transplant, AML027 showed the highest level of blast cells and immature erythroids (CD34+, GATA1+), consistent with the relapse diagnosis and return of the malignant host AML (**Fig. 5c**). These observations would have been difficult to make with FACS analysis, with limited number of markers for early erythroid lineages. Second, ∼20% cells in the AML027 post-transplant sample show markers of immature granulocytes (AZU1, IL8, **Fig. 5c,** **Supplemental Fig. 10d-f**), which are absent in AML035 post-transplant sample, and generally low among AML patients^31^. These cells lack marker expression for mature cells, suggesting the presence of residual precursor cells that may be part of the leukemic clone. Third, monocytes are abundant in both AML patients before transplant (10% and 25% in AML027 and AML035 respectively), but are not detectable after transplant (**Fig. 5c**). Monocytes have been previously identified in post-transplant samples, and the unexpected monocytopenia needs to be followed up with additional studies. Taken together, the analysis provided insights into the cellular composition and possible presence of residual disease in the bone marrows of HSCT recipients that was not available from routine clinical assays.

## DISCUSSION

Here we present a droplet-based scRNA-seq technology that enables digital profiling of thousands to tens of thousands of cells per sample. We demonstrated its application in profiling large immune systems, where we studied the substructures within 68k PBMCs. The ability of GemCode technology to generate faithful scRNA-seq profiles from cryopreserved samples with high cell capture efficiency enabled the application of the scRNA-seq analysis to clinical samples. We successfully generated scRNA-seq samples from fragile BMMCs of transplant samples, and correctly estimated the proportion of donor and host genotypes. In addition, our clustering analysis provided a richer understanding of the complex interplay between host and donor cells and of multiple lineages in the post-transplant setting. It sheds light on the early erythroid lineage in patients, and offered a more comprehensive assessment of patients’ disease progression than would have been possible with routine FACS analysis and clinical chimerism tests. It is our belief that the GemCode single cell technology will in the near term greatly expand research possibilities for clinicians and basic scientists, and will ultimately lead to improved patient care.

### Accession codes

Single cell RNA-seq data have been deposited in the Short Read Archive (SRA) under accession number SRP073767. Data is also available at http://support.10xgenomics.com/single-cell/datasets. The analysis code for the 68k PBMC analysis is available at https://github.com/10XGenomics/single-cell-3prime-paper.

## AUTHOR CONTRIBUTIONS

G.X.Y.Z., J.M.T., P.B., P.R., Z.W.B., T.S.M., B.J.H., J.H.B., E.A.S., and J.P.R. designed experiments. J.M.T., P.B., Z.W.B., S.B.Z., T.D.W., G.P.M., J.S., L.M., S.Y.N., E.A.S., N.G.E., L.W.B., H.J.D., C.M., K.R.L., and W.J.V. conducted experiments. T.D.W., D.A.M., R.B., K.D.N., and B.J.H. designed the instrument. G.P.M., Z.W.B., S.Y.N., C.M.H., P.W.W., and K.D.N. designed the reagents. P.R., R.W., A.W., G.X.Y.Z., J.J.Z., T.S.M., and M.S.L. wrote the analysis software. G.X.Y.Z., P.R., T.S.M., J.Z., K.R.L., and M.T.G. analyzed the data. G.X.Y.Z., E.A.S., J.P.R., T.S.M., B.J.H., and J.H.B. wrote the manuscript.

## ACKNOWLEDGEMENTS

We thank Deanna Church for critical reading of the manuscript, and members of the Bielas laboratory and 10x Genomics for helpful discussions. We thank members of the clinical immunogenetics laboratory at the Fred Hutchinson Cancer Research Center for their assistance in sample preparation and data review: David Wu, Debra Cordell, Aida Guzman, Reena Patel, Ada Ng, Chuck Kellum, and Gana Balgansuren. We acknowledge support from the Canary Foundation (to J.H.B.), an Ellison Medical Foundation New Scholar Award (AG-NS-0577-09 to J.H.B.), an Outstanding New Environmental Scientist Award (ONES) (R01) from the National Institute of Environmental Health Sciences (R01ES019319 to J.H.B.), a grant from the Congressionally Directed Medical Research Programs/U.S. Department of Defense (W81XWH-10-1-0563 to J.H.B.), the Pacific Ovarian Cancer Research Consortium Ovarian Cancer SPORE Award (P50 CA083636). W.J.V is supported by an Achievement Rewards for College Scientists (ARCS) Foundation Fellowship, and Ruth L. Kirschstein National Research service F30 Award for Individual Predoctoral MD/PhD Degree Fellows (NCI F30CA200247).

## COMPETING INTERESTS

G.X.Y.Z., J.M.T., P.B., P.R., Z.W.B., R.W., S.B.Z., T.D.W., J.J.Z., G.P.M., J.S., L.M., D.A.M., S.Y.N., M.S.L., P.W.W., C.M.H., R.B., A.W., K.D.N., T.S.M., and B.J.H. are employees of 10x Genomics.

## ONLINE METHODS

### High speed imaging of gel beads and cells in GEMs

A microscope (Nikon Ti-E, 10X objective) and a high speed video camera (Photron SA5, frame rate=4000/s) were used to image every GEM as they were generated in the microfluidic chip. A custom analysis software was used to count the number of GEMs generated and the number of beads present in each GEM, based on edge detection and the contrast between bead edges and GEM edges and the adjacent liquid. The results of the analysis are summarized in **Fig. 1c**. To estimate the distribution of cells in GEMs, manual counting was used for ∼28k frames of one video on a subset of GEMs. The results indicate an approximate adherence to a Poisson distribution. However, the percentage of multiple cell encapsulations was 16% higher than the expected value, possibly due to sub-sampling error or to cell-cell interactions (some two-cell clumps were observed during the manual count) (**Supplementary Fig. 1b**).

### Cell lines and transplant patient samples

Jurkat (ATCC TIB-152), 293T (ATCC CRL-11268) and 3T3 (ATCC CRL-1658) cells were acquired from ATCC and cultured according to ATCC guidelines. Fresh PBMCs, frozen PBMCs and BMMCs were purchased from ALLCELLS.

The Institutional Review Board at the Fred Hutchinson Cancer Research Center approved the study on transplant samples. The procedures followed were in accordance with the Helsinki Declaration of 1975 and the Common Rule. Samples were obtained after patients had provided written informed consent on molecular analyses. We identified patients with AML undergoing allogeneic hematopoietic stem cell transplant at the Fred Hutchinson Cancer Research Center. The diagnosis of AML was established according to the revised criteria of the World Health Organization^32^.

Bone marrow aspirates were obtained for standard clinical testing 20-30 days before transplant and serially post-transplant according to the treatment protocol. Bone marrow aspirate aliquots were processed within 2 hours of the draw. The BMMCs were isolated using centrifugation through a Ficoll gradient (Histopaque-1077, Sigma Life Science, St Louis, MO). The BMMCs were collected from the serum-Ficoll interface with a disposable Pasteur pipet and transferred to the 50ml conical tube with 2% patient serum in 1xPBS. The BMMCs were counted using a hemacytometer and viability was assessed using Trypan Blue. The BMMCs were resuspended in 90% FBS, 10% DMSO freezing media and frozen using a Thermo Scientific Nalgene Mr. Frosty (Thermo Scientific) in a −80°C freezer for 24 hours before transferred to liquid nitrogen for long-term storage.

### Estimation of RNA content per cell

The amount of RNA per cell type was determined by quantifying (Qubit, Invitrogen) RNA extracted (Maxwell RSC simplyRNA Cells Kit) from several different known number of cells.

### Cell preparation

Fresh cells were harvested, washed with 1x PBS and resuspended at 1×10^6^ cells/ml in 1x PBS and 0.04% BSA. Fresh PBMCs were frozen at 10x by resuspending PBMCs in DMEM + 40% FBS + 10% DMSO, freezing to −80°C in a CoolCell® FTS30 (BioCision), then placed in liquid nitrogen for storage.

Frozen cell vials from ALLCELLS and transplant studies were rapidly thawed in a 37°C water bath for approximately 2 minutes. Vials were removed when a tiny ice crystal was left. Thawed PBMCs were washed twice in medium then resuspended in 1x PBS and 0.04% BSA at room temperature. Cells were centrifuged at 300 rcf for 5 min each time. Thawed BMMCs were washed and resuspended in 1x PBS and 20% FBS. The final concentration of thawed cells was 1×10^6^ cells/ml.

### Sequencing library construction using the GemCode platform

Cellular suspensions were loaded on a GemCode Single Cell Instrument (10x Genomics, Pleasanton, CA) to generate single cell GEMs. Single cell RNA-Seq libraries were prepared using GemCode Single Cell 3’ Gel Bead and Library Kit (now sold as P/N 120230, 120231, 120232, 10x Genomics). GEM-RT was performed in a C1000 Touch™ Thermal cycler with 96-Deep Well Reaction Module (Bio-Rad P/N 1851197): 55°C for 2 hours, 85°C for 5 minutes; held at 4°C. After RT, GEMs were broken and the single strand cDNA was cleaned up with DynaBeads® MyOne™ Silane Beads (Thermo Fisher Scientific P/N 37002D) and SPRIselect Reagent Kit (0.6X SPRI, Beckman Coulter P/N B23318). cDNA was amplified using the C1000 Touch™ Thermal cycler with 96-Deep Well Reaction Module: 98°C for 3 min; cycled 14x: 98°C for 15s, 67°C for 20s, and 72°C for 1 min; 72°C for 1 min; held at 4°C. Amplified cDNA product was cleaned up with the SPRIselect Reagent Kit (0.6X SPRI). The cDNA was subsequently sheared to ∼200bp using a Covaris M220 system (Covaris P/N 500295). Indexed sequencing libraries were constructed using the reagents in the GemCode Single Cell 3’ Library Kit, following these steps: 1) end repair and A-tailing; 2) adapter ligation; 3) post-ligation cleanup with SPRIselect; 4) sample index PCR and cleanup. The barcode sequencing libraries were quantified by quantitative PCR (qPCR) (KAPA Biosystems Library Quantification Kit for Illumina platforms P/N KK4824). Sequencing libraries were loaded at 2.1pM on an Illumina NextSeq500 with 2 × 75 paired-end kits using the following read length: 98bp Read1, 14bp I7 Index, 8bp I5 Index and 10bp Read2. Some earlier libraries were made with 5nt UMI, and 5bp Read2 was obtained instead. These libraries have been documented in **Supplementary Table 2**.

### ERCC assay

ERCC synthetic spike-in RNAs (Thermo Fisher P/N 4456740) were diluted (1:10 or 1:50) and loaded into a GemCode Single Cell Instrument, replacing cells normally used to generate GEMs. Spike-in Mix1 and Mix2 were both tested. A slightly modified protocol was used as only a small fraction of GEMs were collected for RT and cDNA amplification. After the completion of GEM-RT, 1.25 µL of the emulsion was removed and added to a bi-phasic mixture of Recovery Agent (125 µL) (P/N 220016) and 25 mM Additive 1 (30 µL) (P/N 220074, 10x Genomics). The recovery agent was then removed and the remaining aqueous solution was cleaned up with the SPRISelect Reagent Kit (0.8X SPRI). cDNA was amplified using the C1000 Touch™ Thermal cycler with 96-Deep Well Reaction Module: 98°C for 3 min; cycled 14x: 98°C for 15s, 67°C for 20s, and 72°C for 1 min; 72°C for 1 min; held at 4°C. Amplified cDNA product was cleaned up with the SPRIselect Reagent Kit (0.8X) cDNA was subsequently sheared to ∼200bp using a Covaris M220 system to construct sample-indexed libraries with 10x Genomics adapters. Expected ERCC molecule counts were calculated based on the amount of ERCC molecules used and sample dilution factors. The counts were compared to detected molecule counts (UMI counts) to calculate conversion efficiency.

### ddPCR assay

Jurkat cells were used in ddPCR assays to estimate conversion efficiency as follows. 1) The amount of RNA per Jurkat cell was determined by quantifying (Qubit, Invitrogen) RNA extracted (Maxwell RNA Purification Kits) from several different known number of Jurkat cells. 2) Bulk RT-ddPCR (Bio-Rad One-Step RT-ddPCR Advanced Kit for Probes 1864021) was performed on the extracted RNA to determine the copy number per cell of 8 selected genes. 3) Approximately 5000 Jurkat cells were processed using the GemCode Single Cell 3’ platform, and single stranded cDNA was collected after RT in GEMs following the protocols listed in “Sequencing library construction using the GemCode platform”. cDNA copies of the 8 genes were determined using ddPCR (Bio-Rad ddPCR Supermix for Probes (no dUTP) P/N 1863024). The actual Jurkat cell count was found by sequencing a subset of the GEM-RT reactions on a MiSeq. The conversion efficiency is the ratio between cDNA copies per cell (step 3) and RNA copies per cell from bulk RT-ddPCR (step 2), assuming a 50% efficiency in RT-ddPCR^33^.

The probe sequences for the ddPCR assay are as follows.

SERAC1_f: CACGAGCCGCCAGC; SERAC1_r: TCTGCAACAGATGACGCAATAAG;

SERAC1_p: /56-FAM/CGCCTGCCG/ZEN/GCAGAATGTC/3IABkFQ/.

AP1S3_f: GAAGCAGCCATGGTCTAAGC; AP1S3_r: CCTTGTCGACTGAAGAGCAATATG;

AP1S3_p: /56-FAM/CGGCCCAGC/ZEN/CACGATGATACAT/3IABkFQ/OR.

AOV1_f: CCGGAAGTGGGTCTCGTOR; AOV1_r: TTCTTCATAGCCTTCCCGATACCOR;

AOV1_p: /56-FAM/TCGTGATGG/ZEN/CGGATGAGAGGTTTCA/3IABkFQ/.

DOLPP1_f: ATGGCAGCGGACGGA; DOLPP1_r: GGCTCAGGTAGGCAAGGA; DOLPP1_p: /56-FAM/CCACGTCGA/ZEN/ATATCCTGCAGGTGATCT/3IABkFQ/.

KPNA6_f: TGAAAGCTGCCGCTGAAG; KPNA6_r: CCCTGGGCTCGCCAT; KPNA6_p: /56-FAM/CGGACCCGC/ZEN/GATGGAGACC/3IABkFQ/.

ITSN2_f: GTGACAGGCTACGCAACAG; ITSN2_r: TCCTGAGTTTTCCTTGCTAGCT; ITSN2_p: /56-FAM/AGGGCGCCA/ZEN/GATGGCTGA/3IABkFQ/.

LCMT1_f: GTCGACCCCGCTTCCA; LCMT1_r: GGTCATGCCAGTAGCCAATG; LCMT1_p: /56-FAM/ATGCTTCCC/ZEN/TGTGCAAGAGGTTTGC/3IABkFQ/.

AP2M1_f: GCAGCGGGCAGACG; AP2M1_r: ATGGCGGCAGATCAGTCT; AP2M1_p: /56-FAM/CATCGCTCT/ZEN/GAGAACAGACCTGGTG/3IABkFQ/.

### Cell capture efficiency calculation

The efficiency is calculated by taking the ratio of the number of cells detected by sequencing vs. the number of cells loaded into the chip. The latter is determined from (volume added*input concentration of cells), and takes into account losses in the chip. These losses include: 1) cells left behind in sample well, 2) cells in GEMs left behind in the outlet well, 3) cells in GEMs with Nbead = 0 and Nbead > 1. The losses do not include cells left behind in pipette tips during mixing and transfer steps before pipetting into the sample well. The theoretical efficiency (based on the Cell Loading Correction Factor of 1.92) is 52%. It is worth noting that there is a 15-20% error in cell counts, which could account for at least some of the variability in the calculated efficiencies.

### Chimerism assay

PowerPlex 16 System (Promega) was used in conjunction with an Applied Biosystems (Life Technologies) 3130xl Genetic Analyzer. Donor BMMCs were used as the reference baseline.

### Alignment, barcode assignment and UMI counting

The Cell Ranger Single Cell Software Suite was used to perform sample demultiplexing, barcode processing, and single cell 3’ gene counting (http://software.10xgenomics.com/single-cell/overview/welcome). First, sample demultiplexing was performed based on the 8bp sample index read to generate FASTQs for the Read1 and Read2 paired-end reads as well as the 14bp GemCode barcode. 10bp UMI tags were extracted from Read2. Then, Read1, which contains the cDNA insert, was aligned to an appropriate reference genome using STAR^34^. For mouse cells, mm10 was used. For human cells, hg19 was used. For samples with mouse and human cell mixtures, the union of hg19 and mm10 were used. For ERCC samples, ERCC reference (https://tools.thermofisher.com/content/sfs/manuals/cms_095047.txt) was used.

Next, GemCode barcodes and UMIs were filtered. All of the known listed of barcodes that are 1-Hamming-distance away from an observed barcode are considered. Then the posterior probability that the observed barcode was produced by a sequencing error is computed, given the base qualities of the observed barcode and the prior probability of observing the candidate barcode (taken from the overall barcode count distribution). If the posterior probability for any candidate barcode is at least 0.975, then the barcode is corrected to the candidate barcode with the highest posterior probability. If all candidate sequences are equally probable, then the one appearing first by lexical order is picked.

UMIs with sequencing quality score>10 were considered valid if they were not homopolymers. Qual=10 implies 90% base call accuracy. A UMI that is 1-Hamming-distance away from another UMI (with more reads) for the same cell barcode and gene, is corrected to the UMI with more reads. This approach is nearly identical to that in Jaitin *et al*.^4^, and is similar to that in Klein *et al.*^8^ (although Klein *et al.*^8^ also used UMIs to resolve multi-mapped reads, which was not implemented here).

Lastly, PCR duplicates were marked if two sets of read pairs shared a barcode sequence, a UMI tag, and a gene ID (Ensembl GTFs GRCh37.82, ftp://ftp.ensembl.org/pub/grch37/release-84/gtf/homo_sapiens/Homo_sapiens.GRCh37.82.gtf.gz, and GRCm38.84, ftp://ftp.ensembl.org/pub/release-84/gtf/mus_musculus/Mus_musculus.GRCm38.84.gtf.gz, were used). Only confidently mapped (MAPQ=255), non-PCR duplicates with valid barcodes and UMIs were used to generate gene-barcode matrix.

Cell barcodes were determined based on distribution of UMI counts. All top barcodes within the same order of magnitude (greater than 10% of the top nth barcode where n is 1% of the expected recovered cell count) were considered cell barcodes. Number of reads that provide meaningful information is calculated as the product of 4 metrics: 1) valid barcodes; 2) valid UMI; 3) associated with a cell barcode; and 4) confidently mapped to exons.

In the mouse and human mixing experiments, multiplet rate was defined as twice the rate of cell barcodes with significant UMI counts from both mouse and human, where top 1% of UMI counts was considered significant. The extent of barcode crosstalk was assessed by the fraction of mouse reads in human barcodes, or vice versa.

Samples processed from multiple channels can be combined by concatenating gene-cell-barcode matrices. This functionality is provided in the Cell Ranger R Kit. Sequencing data from multiple sequencing runs of a library can be combined by counting non-duplicated reads. This functionality is provided in the Cell Ranger pipeline. In addition, sequencing data can be subsampled to obtain a given number of UMI counts per cell. This functionality is also provided in the Cell Ranger R Kit, and is useful when combining data from multiple samples for comparison.

### PCA analysis of mixing of Jurkat and 293T cells

Gene-cell-barcode matrix from each of the 4 samples was concatenated. Only genes with at least 1 UMI count detected in at least 1 cell are used. UMI normalization was performed by first dividing UMI counts by the total UMI counts in each cell, followed by multiplication with the median of the total UMI counts across cells. Then we took the natural log of the UMI counts. Finally, each gene was normalized such that the mean signal for each gene is 0, and standard deviation is 1. PCA was run on the normalized gene-barcode matrix. The normalized UMI counts of each gene is used to show expression of a marker in a tSNE plot.

### SNV analysis of Jurkat and 293T scRNA-seq data

SNVs were called by running Freebayes 1.0.2^35^ on the genome BAM produced by Cell Ranger. High quality SNVs (SNV calling Qual>=100 with at least 10 UMI counts from at least 2 cells; indels are ignored) that were only observed in Jurkat or 293T cells (but not both) were selected. Cells were labeled as Jurkat or 293T based on Jurkat- and 293T-specific SNV counts, where the fraction of counts from the other species is <0.2. Cells with fraction of SNV from either species between 0.2 and 0.8 are considered multiplets. The inferred multiplet rate is 2* observed multiplet rate (to account for Jurkat:Jurkat and 293T:293T multiplets).

### PCA and t-SNE analysis of PBMCs

Genes with at least 1 UMI count detected in at least 1 cell are used. Top 1000 most variable genes were identified based on their mean and dispersion (variance/mean), which is similar to the approach used by Macoscko *et al*^7^. Genes were placed into 20 bins based on their mean expression. Normalized dispersion is calculated as the absolute difference between dispersion and median dispersion of the expression mean, normalized by median absolute deviation within each bin.

PCA was run on the normalized gene-barcode matrix of the top 1000 most variable genes to reduce the number of feature (gene) dimensions. UMI normalization was performed by first dividing UMI counts by the total UMI counts in each cell, followed by multiplication with the median of the total UMI counts across cells. Then we took the natural log of the UMI counts. Finally, each gene was normalized such that the mean signal for each gene is 0, and standard deviation is 1. PCA was run on the normalized gene-barcode matrix. After running PCA, Barnes-hut^36^ approximation to t-distributed Stochastic Neighbor Embedding (t-SNE)^16^ was performed on the first 50 PCs to visualize cells in a 2-D space. 50 PCs were used because: 1) using all PCs would take a very long time with tSNE analysis; 2) they explained ∼25% of total variance. K-means^15^ clustering was run to group cells for the clustering analysis. k=10 was selected based on the sum of squared error scree plot (**Supplementary Fig. 5d**).

### Identification of cluster-specific genes and marker-based classification

To identify genes that are enriched in a specific cluster, the mean expression of each gene was calculated across all cells in the cluster. Then each gene from the cluster was compared to the median expression of the same gene from cells in all other clusters. Genes were ranked based on their expression difference, and the top 10 enriched genes from each cluster were selected. For hierarchical clustering, pair-wise correlation between each cluster was calculated, and centered expression of each gene was used for visualization by heatmap.

Classification of PBMCs was inferred from the annotation of cluster-specific genes. In the case of cluster 10, marker expression of multiple cell types (e.g. B, dendritic, and T) was detected. Since the relative cluster size of B, dendritic and T is 5.7%, 6.6% and 81% respectively, we’d expect the cluster 10 (which is only 0.5%) to contain multiplets consisting mostly from B:dendritic (0.36%) and B:dendritic:T (0.3%).

### Selection of purified sub-populations of PBMCs

Each population of purified PBMCs was downsampled to ∼16k reads per cell. PCA, tSNE and k-means clustering were performed for each downsampled matrix, following the same steps outlined in **PCA and t-SNE analysis of PBMCs**. Only one cluster was detected in most samples, consistent with the FACS analyses (**Supplementary Fig. 6**). For samples with more than one cluster, only clusters that displayed the expected marker gene expression were selected for downstream analysis. For CD14+ Monocytes, 2 clusters were observed and identified as CD14+ Monocytes and Dendritic cells based on expression of marker genes FTL and CLEC9A, respectively.

### Cell classification analysis using purified PBMCs

Each population of purified PBMCs was downsampled to ∼16k confidently mapped reads per cell. Then, an average (mean) gene expression profile across all cells was calculated. Next, gene expression from every cell of the complex population was compared to the gene expression profiles of purified populations of PBMCs by spearman correlation. The cell was assigned the ID of the purified population if it had the highest correlation with that population. Note that the difference between the highest and 2^nd^ highest correlation was small for some cells (for example, the difference between cytotoxic T and NK cells), suggesting that the cell assignment was not as confident for these cells. A few of the purified PBMC populations overlapped with each other. For example, CD4+ T Helper 2 cells include all CD4+ cells. This means that cells from this sample will overlap with cells from samples that contain CD4+ cells, including CD4+/CD25+ T Reg, CD4+/CD45RO+ T Memory, CD4+/CD45RA+/CD25-Naïve T. Thus, when a cell was assigned the ID of CD4+ T Helper 2 cell based on the correlation score, the next highest correlation was checked to see if it was one of the CD4+ samples. If it was, the cell’s ID was updated to the cell type with the next highest correlation. The same procedure was performed for CD8+ Cytotoxic T and CD8+/CD45RA+ Naïve Cytotoxic T (which is a subset of CD8+ Cytotoxic T).

The R code used to analyze 68k PBMCs and purified PBMCs can be found here: https://github.com/10XGenomics/single-cell-3prime-paper.

### Cell clustering and classification with Seurat

The gene-cell-barcode matrix of 68k PBMCs was log-transformed as an input to Seurat. The top 469 most variable genes selected by Seurat were used to compute the PCs. The first 22 PCs were significant (p < 0.01)based on the built-in jackstraw analysis, and used for tSNE visualization. Cell classification was taken from Cell classification analysis using purified PBMCs.

### Comparison between fresh vs. frozen PBMCs

The sequencing data of 68k fresh PBMCs and 3k frozen PBMCs were down-sampled such that each sample has ∼14k confidently mapped reads/cell. Only genes that are detected in at least one cell were included for the comparison, which uses the mean of each gene across all cells.

For cell classification comparison between purified and frozen PBMCs, we pooled all the cells labeled as T or NK cells together. This is because the sub-populations within T and between T and NK cells are sometimes difficult to cluster separately. We did not want the comparison between fresh vs. frozen cells to be affected by the clustering methods used.

### SNV-based genotype assignment

SNVs were called by running Freebayes 1.0.2^35^ on the genome BAM produced by Cell Ranger. Only SNVs with support from at least 2 cell barcodes, with a minimal SNV Qual score >=30, minimal SNV base Qual>=1 were included. Reference (R) and alternate (A) allele counts were computed at each SNV, producing a matrix of cell-reference UMI counts and cell-alternate-allele UMI counts. These matrices were modeled as a mixture of two genomes where the likelihood of any of the three genotypes (R/R, R/A, or A/A) at a site was taken to be binomially distributed with a fixed error rate of 0.1%. For each sample, two models were inferred in parallel, one where only one genome is present (*K* = 1) and another where two genomes are present (*K* = 2). Inference of the model parameters (cell-to-genome assignments and the *K* sets of genotypes) was performed by using a Gibbs sampler to approximate their posterior distributions. In order to ameliorate the label-switching problem in Monte Carlo inference of mixture models, relabeling of the sampled cell-to-genome assignments was performed as per Stephens *et al*^37^.

In *in silico* cell mixing experiments, when the *K* = 2 model failed to adequately separate the two genomes, it reported a distribution of posterior probabilities near 0.5 for the cell-genome calls, indicating a lack of confidence in those calls. We applied a requirement that 90% of the cells have a posterior probability greater than 75% in order to select the *K* = 2 model over the *K* = 1 model. Selecting *K* = 1 indicates that the mixture fraction is below the level of detection of the method, which in *in silico* mixing experiments was determined to be 4% of 6,000 cells.

### Genotype comparison to the pure sample

To ascertain the assignment of genotypes to individuals, only shared SNVs between the genotype group and the pure sample were considered. Then the average genotype of all the cells was compared to that of the pure sample. In order to obtain some baseline for the % genotype overlap among different individuals, we performed pairwise comparison of genotypes called from the same individuals (11 pairwise comparisons) or from different individuals (15 pairwise comparisons). The percent genotype overlap between the same individuals averages ∼98%±0.3%, whereas the percent genotype overlap between the different individuals averages ∼73%±2%.

### PCA and t-SNE analysis of BMMCs

Data from 6 samples were used: 2 healthy controls, AML027 pre- and post-transplant, and AML035 pre- and post-transplant. Each sample was downsampled to ∼10k confidently mapped reads per cell. Then the gene-cell barcode matrix from each sample was concatenated. PCA, tSNE and k-means clustering were performed on the pooled matrix, following the same steps outlined in **PCA and t-SNE analysis of PBMCs**. For k-means clustering, k=10 was used based on the bend in the sum of squared error scree plot.

Cluster-specific genes were identified following the steps outlined in **Identification of cluster-specific genes and marker-based classification**. Classification was assigned based on cluster-specific genes, and based on expression of some well-known markers of immune cell types. “Blasts and Immature Ery refers to cluster 4, which expresses CD34, a marker of hematopoietic progenitors^38^, and Gata2, a marker for early erythroids^39^. “Immature Ery refers to clusters 5 and 8, which show expression of Gata1, a transcription factor essential for erythropoiesis^40^, but not CD71, which are often found in more committed erythroid cells^38^. “Immature Ery refers to cluster 1, which show expression of CD71. “Mature Ery” refers to cluster 2. HBA1, a marker of mature erythroid cells, is preferentially detected in cluster 2. Cluster 3 was assigned as “Immature Granulocytes” because of the expression of early granulocyte markers such as AZU1 and IL8^41^, and the lack of expression of CD16. Cluster 7 was assigned as “Monocytes” because of the expression of CD14 and FCN1, for example. “B” refers clusters 6 and 9 because of markers such as CD19 and CD79A. “T” refers to cluster 10, because of markers such as CD3D and CD8A.

---

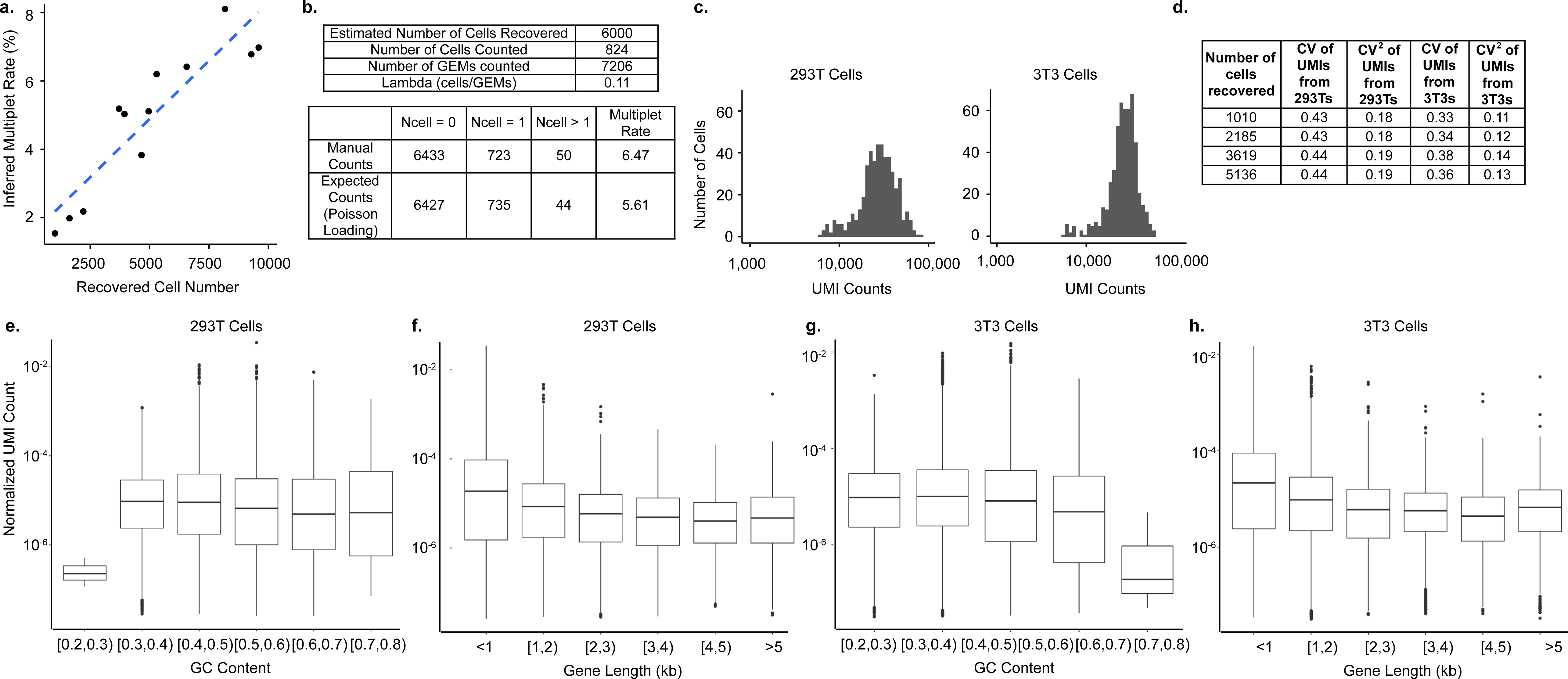

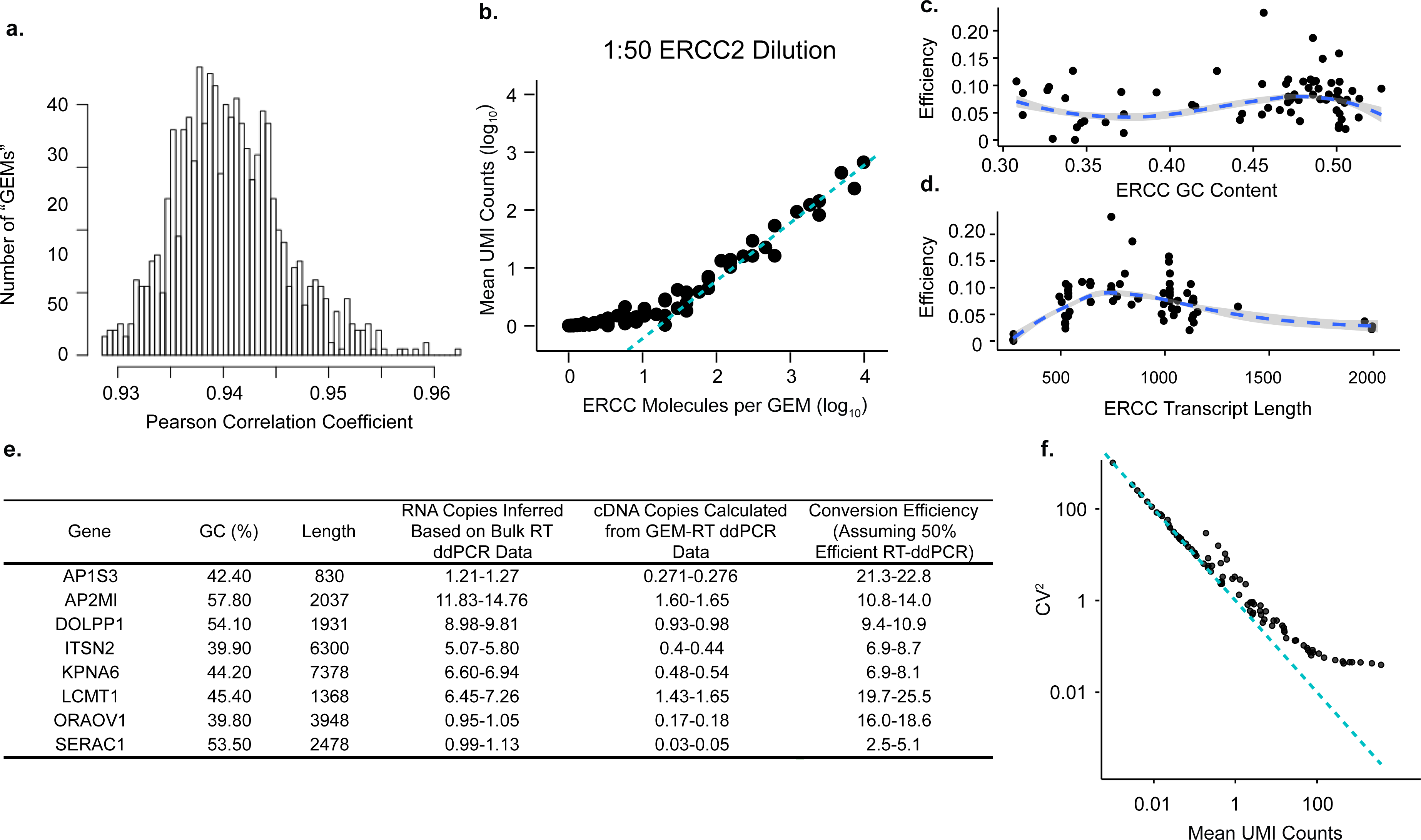

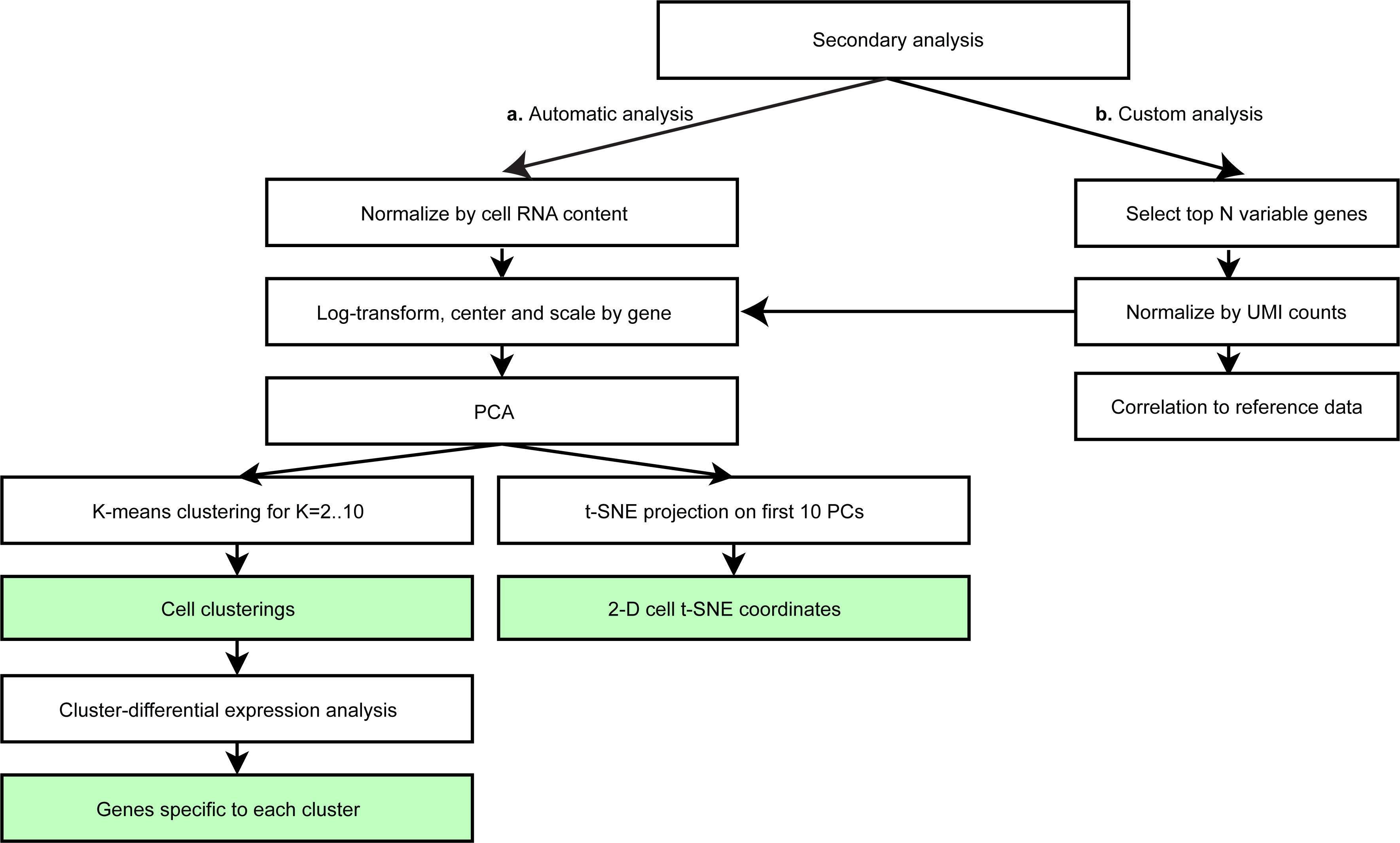

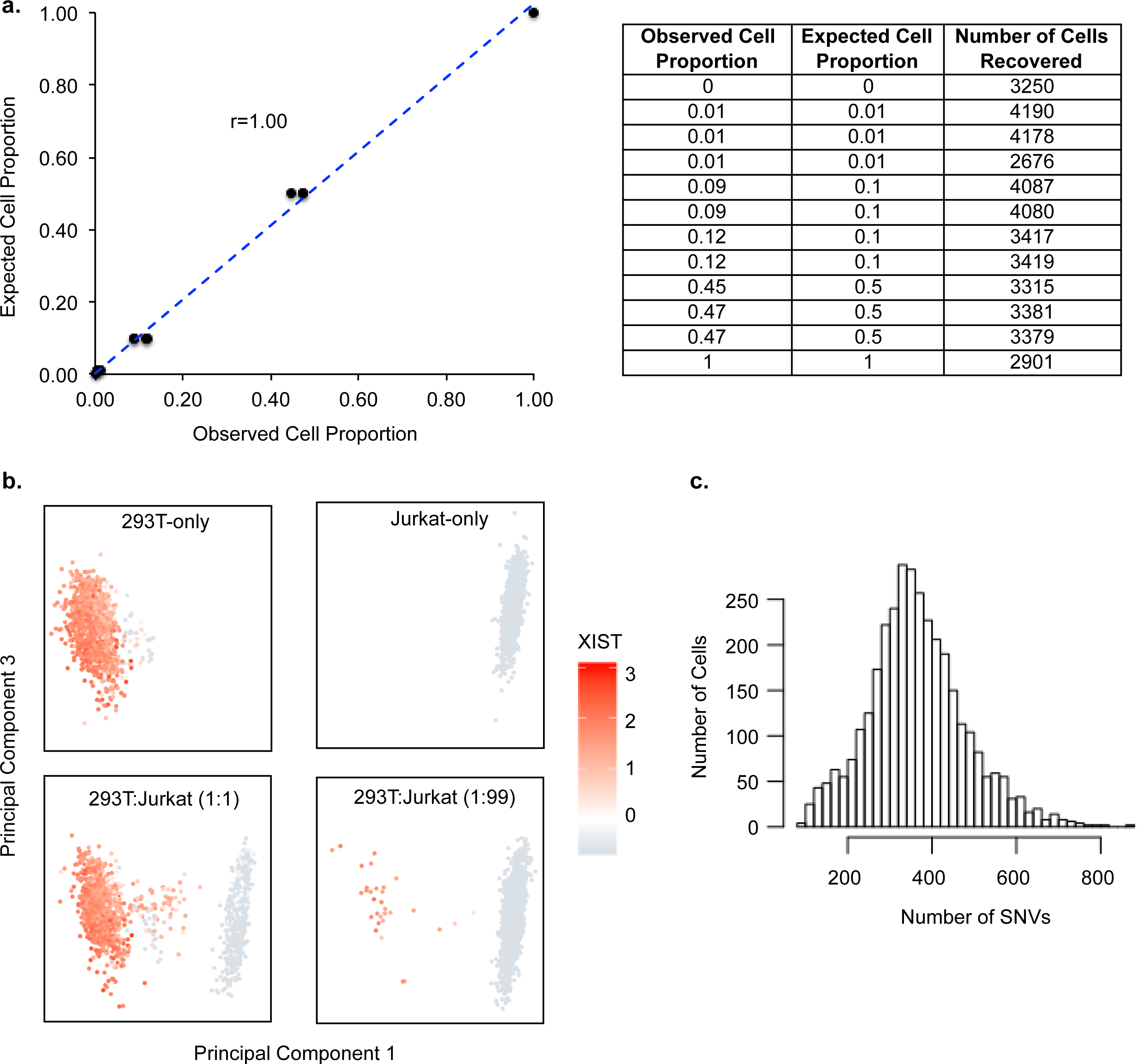

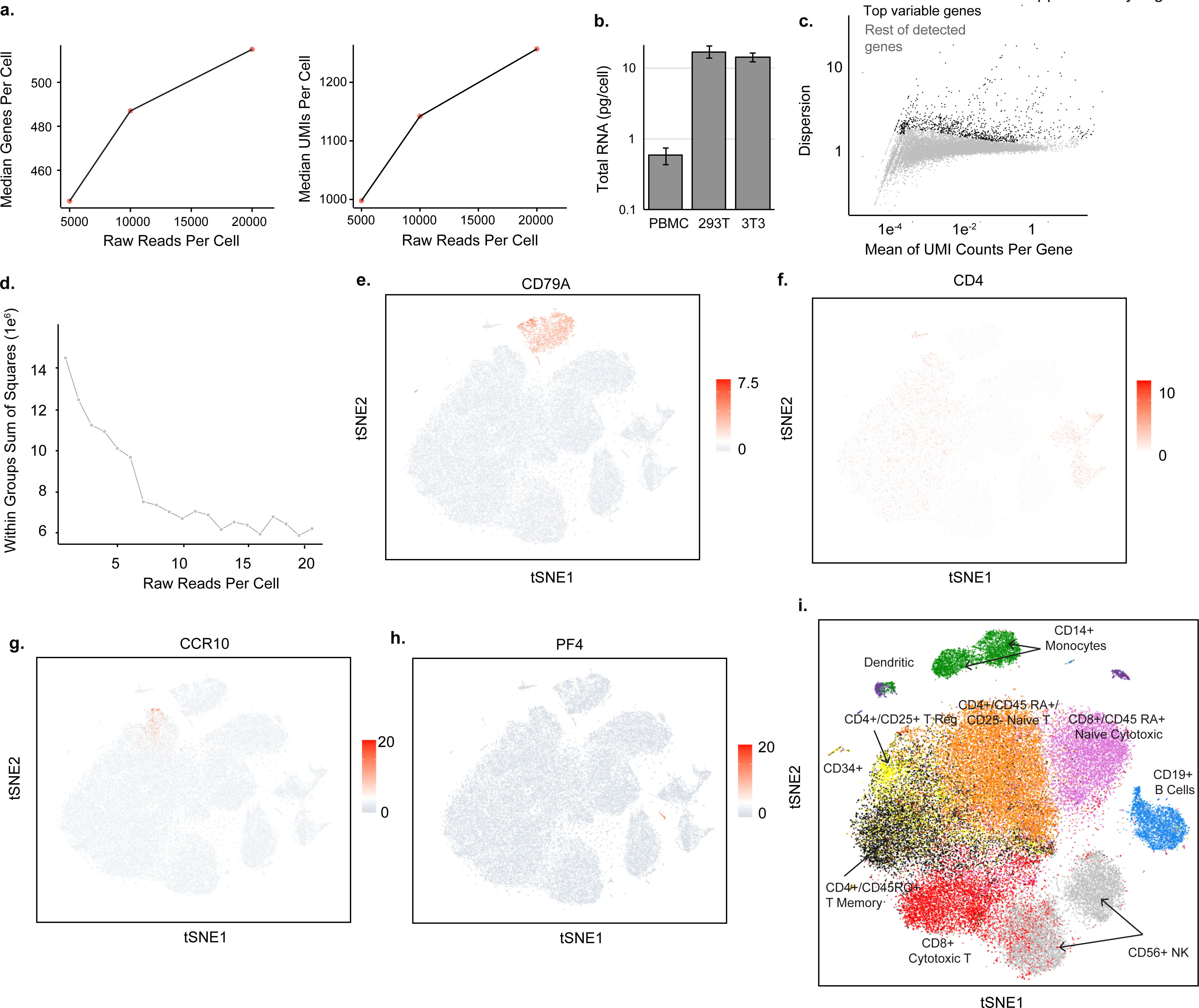

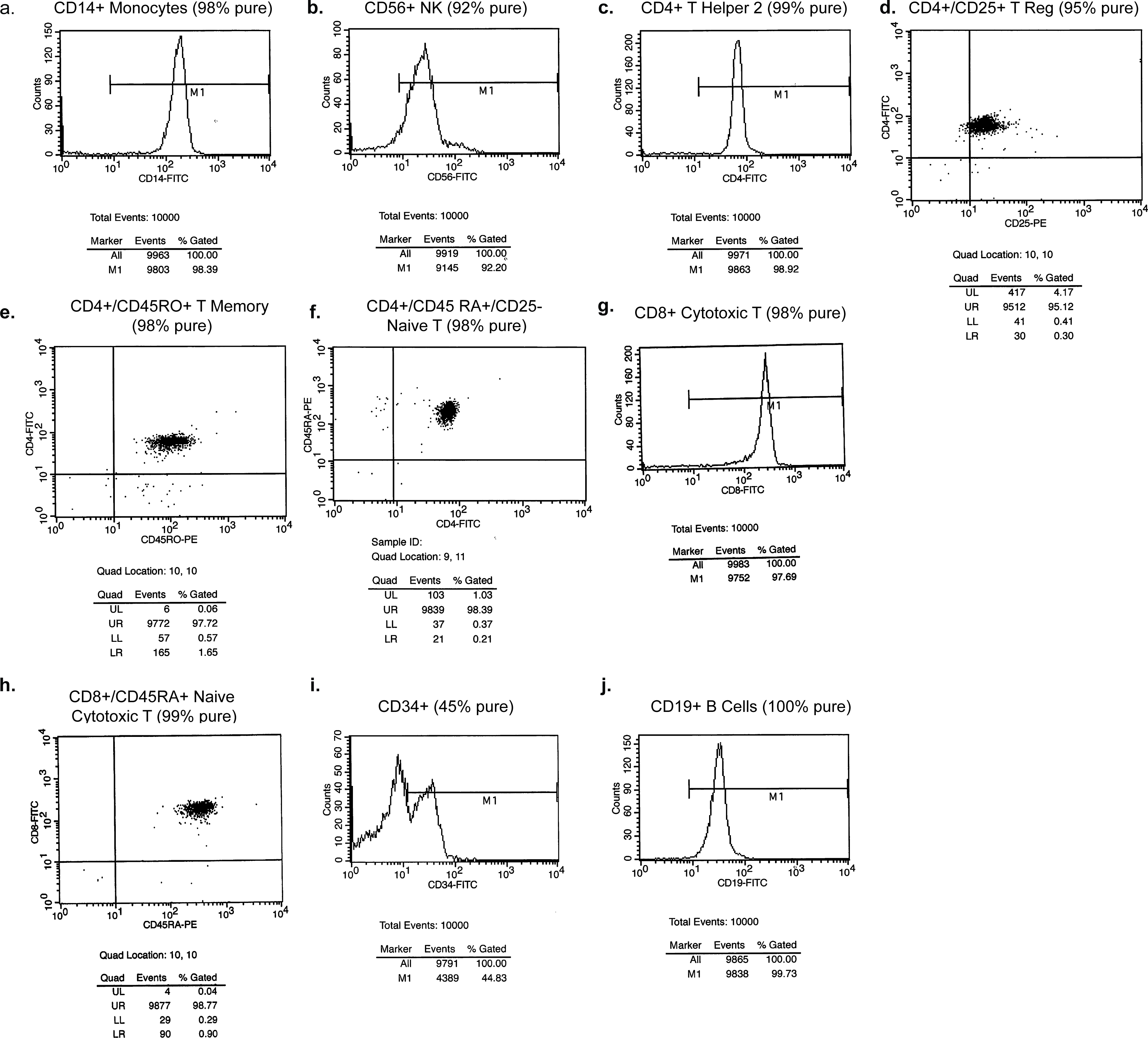

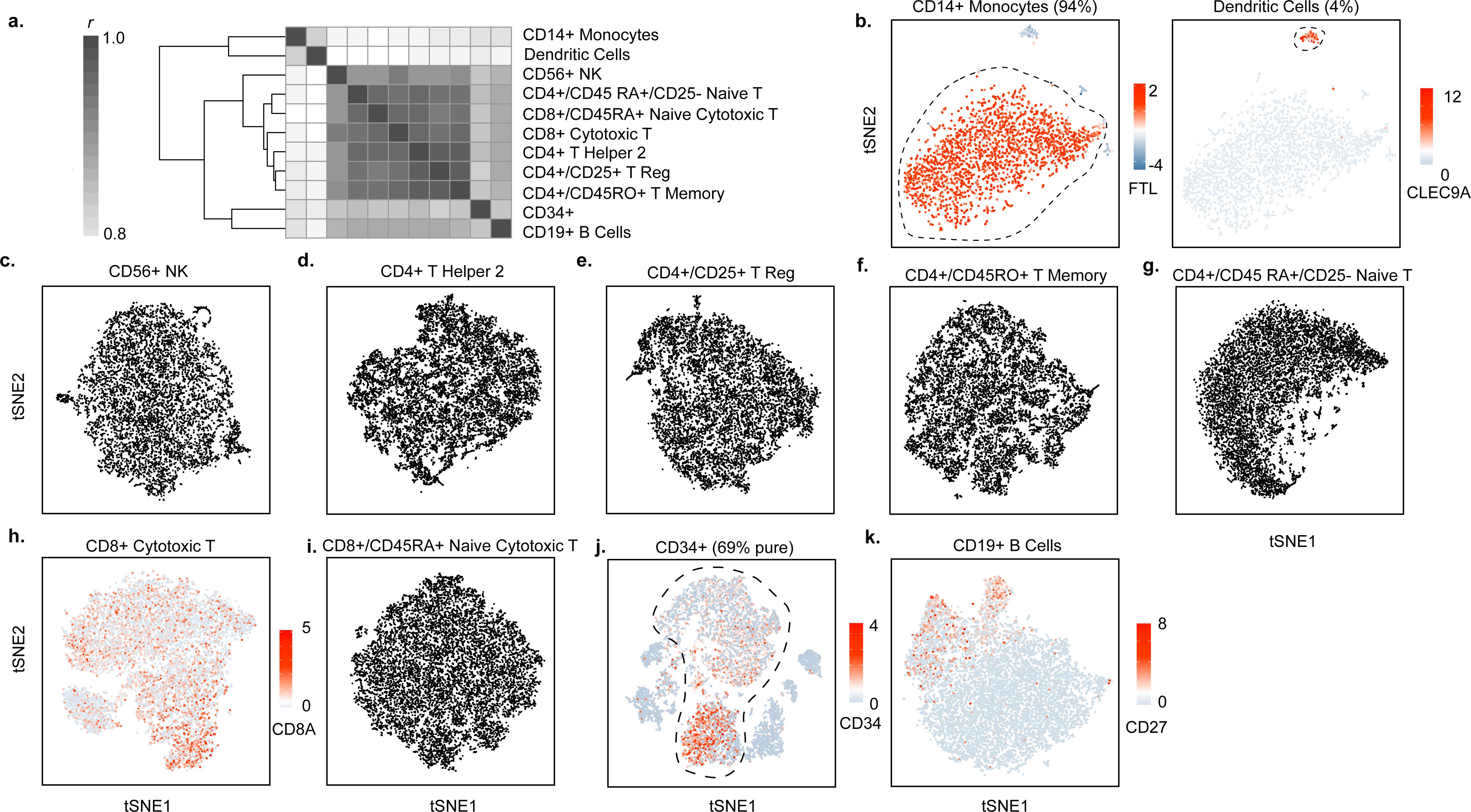

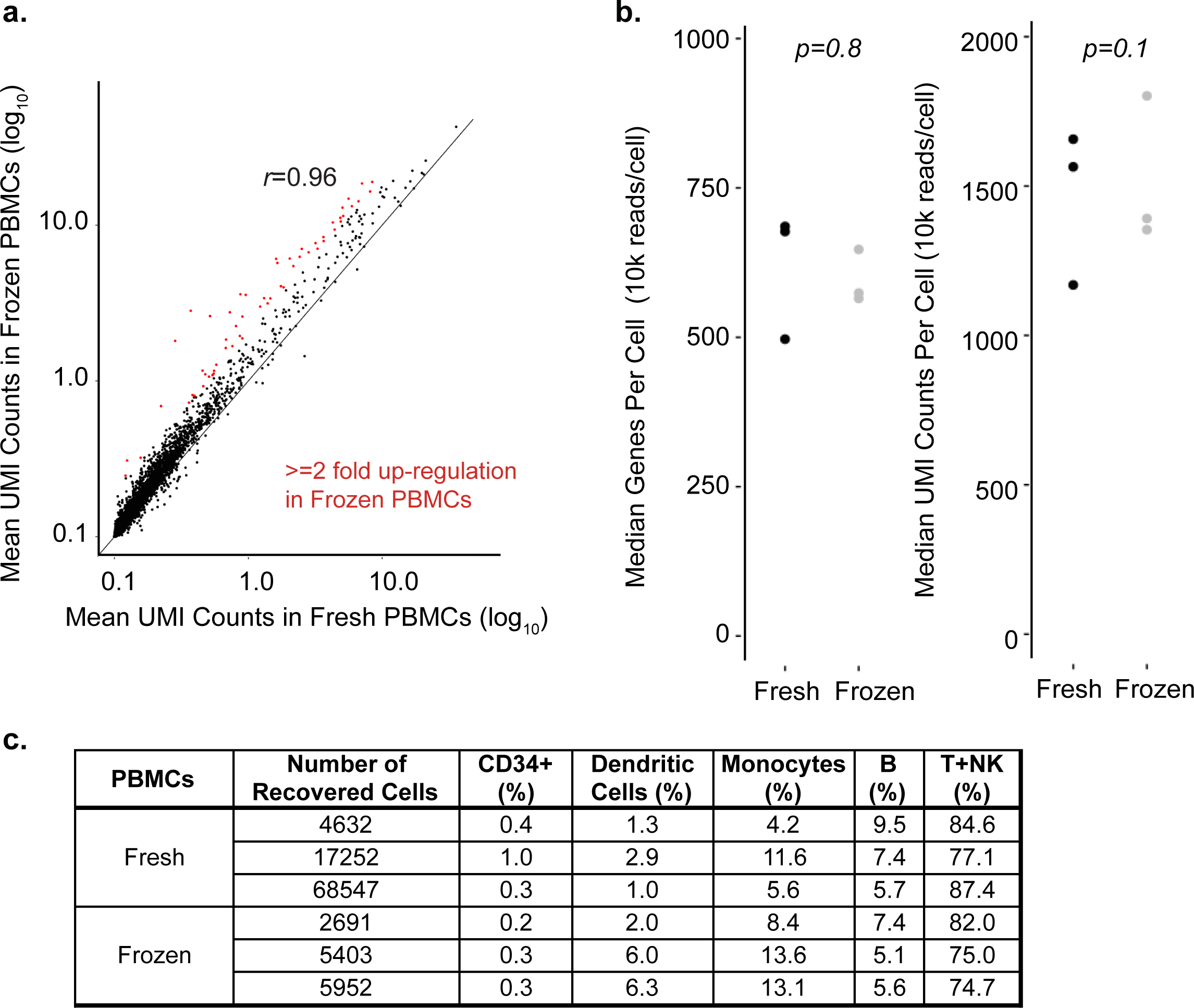

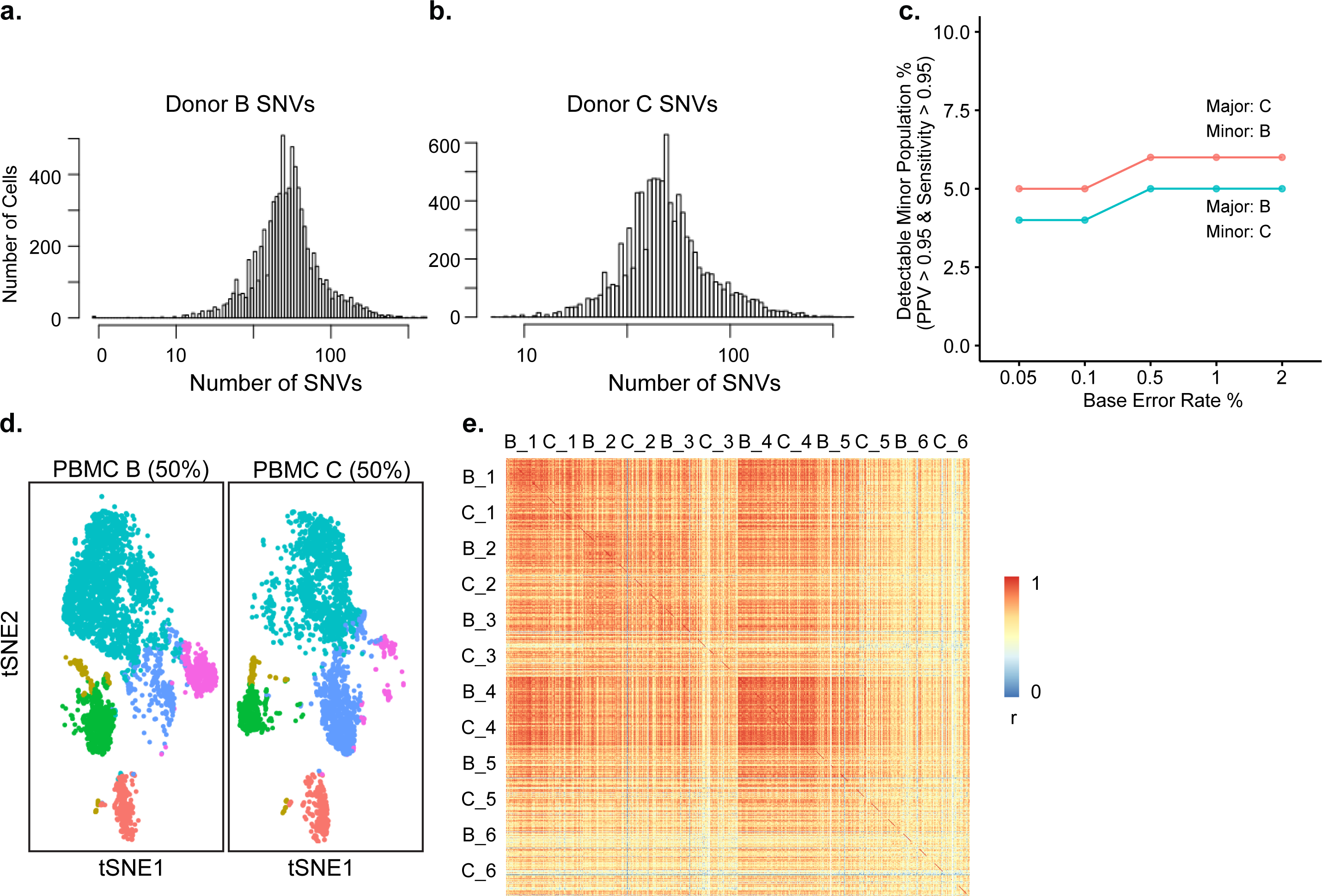

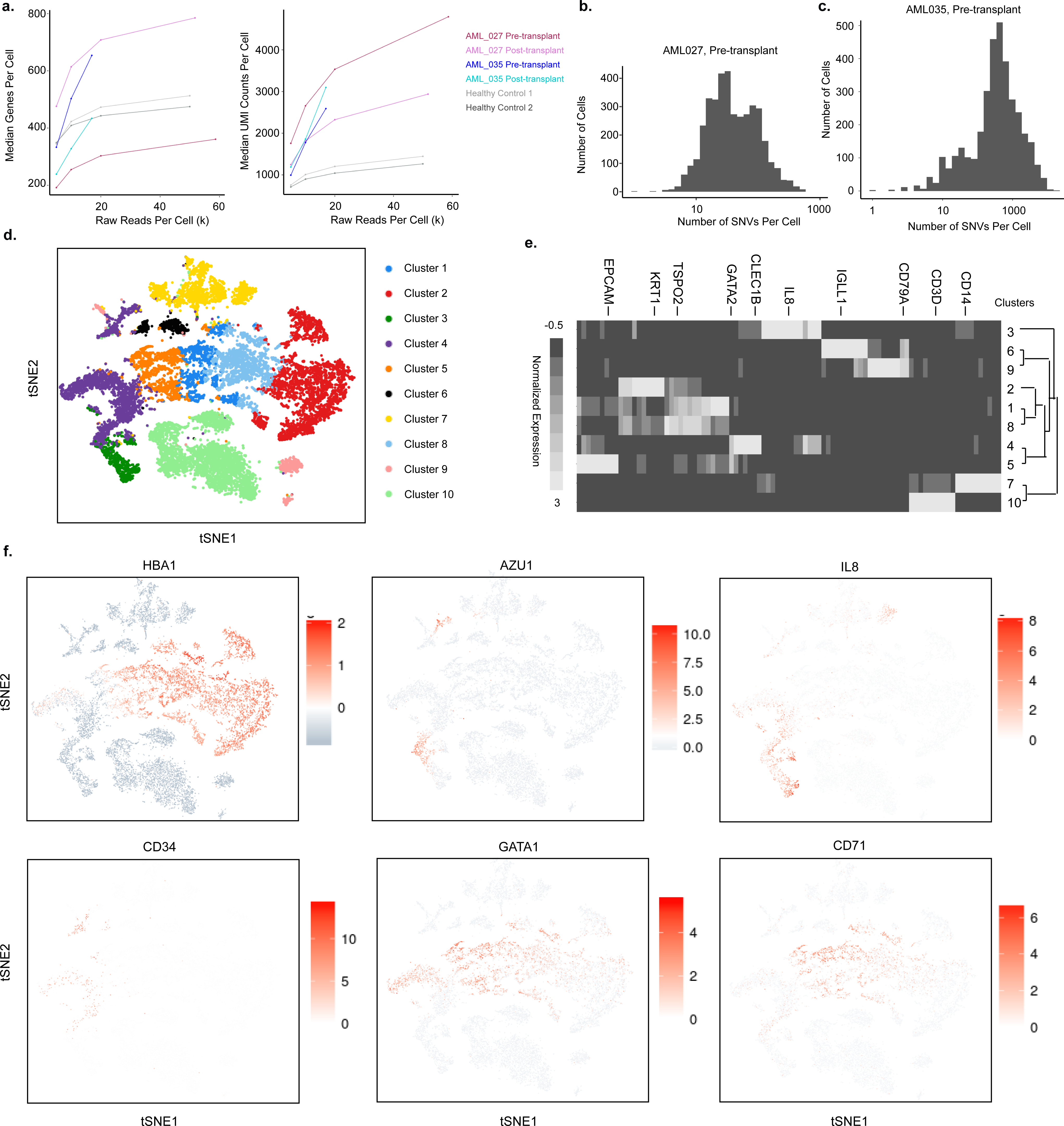

